# Photochromic reversion enables long-term single-molecule tracking in living plants

**DOI:** 10.1101/2024.04.10.585335

**Authors:** Michelle von Arx, Kaltra Xhelilaj, Philip Schulz, Sven zur Oven-Krockhaus, Julien Gronnier

## Abstract

Single-molecule imaging enables the observation of individual molecules in living cells (D’Este et al., 2024; Kusumi et al., 2014; Lelek et al., 2021; Nguyen et al., 2023). In plants, however, the tracking of single molecules is typically limited to a few hundred milliseconds (Bayle et al., 2021; Gronnier et al., 2017; Hosy et al., 2015), precluding the observation of dynamic cellular processes at molecular resolution. Here, we describe photochromic reversion, an imaging modality that enables long-term single-molecule tracking of genetically encoded translational fusions. Using this approach, we achieve minute-long tracking of individual cell-surface receptors and reveal previously inaccessible dynamic spatial arrest events of single plasma membrane proteins. We further developed and benchmarked computational analysis of spatial arrests (CASTA), a machine learning-based tool that automatically detects and analyses spatial, temporal, and diffusional properties of these events, thereby enabling precise nanoscale kinetic measurements. Together, these advances provide a powerful framework for deciphering the principles governing membrane dynamics and function.

## Introduction

Our ability to comprehend fundamental principles of the living is intimately linked to our capacity to observe its molecular intricacy live. Recent progress made in instrumentation and methods permits the detection, imaging, and tracking of single molecules in their native cellular environment (D’Este et al., 2024; Kusumi et al., 2014; Lelek et al., 2021; Nguyen et al., 2023). In animal cells, long-term single-molecule imaging can be achieved by imaging bright and photostable quantum dots (Michalet et al., 2005) or fluorophore ligands bound to SNAP-tag- or HaloTag-tagged proteins of interest (Liu et al., 2018). Due to the limited permeability of walled plant cells, the implementation of similar approaches in plants has been unsuccessful. Single particle photoactivated localization microscopy (spt-PALM) (Manley et al., 2008) has been applied in plants and remains to date the only bona-fide single-molecule tracking (SMT) approach amenable to plant samples (Bayle et al., 2021; Gronnier et al., 2017; Hosy et al., 2015). PALM relies on the iterative and stochastic photo-switching of fluorescent proteins (FP) to spatially and temporally separate single molecules that can be located and tracked with nanoscopic precision. EOS variants are considered the best overall performing photo-switching FP (McKinney et al., 2009; Rodriguez et al., 2017; Wang et al., 2014; Zhang et al., 2012) and are most used for spt-PALM experiments in plants (Bayle et al., 2021; Gronnier et al., 2017; Hosy et al., 2015; Jolivet et al., 2025; Perraki et al., 2018; Platre et al., 2019; Rohr, Ehinger, et al., 2024; Smokvarska et al., 2023; Smokvarska et al., 2020). mEOS variants switch irreversibly from “green” to “red” excitable spectral states upon low intensity illumination at 405 nm. Photo-switched EOS variants have been shown to enter a long-lived dark state which results from chromophore isomerization (De Zitter et al., 2019). Such a long-lived dark state can be reverted to the fluorescent state upon illumination with 488 nm *in vitro* and animal cells (De Zitter et al., 2019). Capitalizing on these observations, we present photochromic reversion, an imaging modality that enables minute-long single-molecule tracking, and Computational Analysis of Spatial Arrests (CASTA), a computational solution for analysing spatial arrest events within single-molecule trajectories. Together, these imaging and computational approaches provide a robust framework for single-molecule studies.

## Results and discussion

We set to investigate the existence of a long-lived dark state of EOS-translational fusions in plants. We imaged translational fusions of the plasma membrane protein REMORIN1.2 (REM1.2) with mEOS2 or mEOS3.2 transiently expressed in *Nicotiana benthamiana* using variable-angle total internal fluorescence microscopy (VA-TIRFM) (Konopka & Bednarek, 2008; Tokunaga et al., 2008). We observed photo-converted EOS molecules without 405 nm light (Supplementary Figure 1A-B) and that illumination of the samples with a low amount of 488 nm light led to the appearance of additional mEOS2-REM1.2 and mEOS3.2-REM1.2 molecules (Supplementary Figure 1A-B) which rapidly disappeared when the 488 nm laser was turned off (Supplementary Figure 1C, Supplementary movie 1). This suggests that mEOS variants enter a long-lived dark state in absence of 488 nm light. To corroborate these observations, we analysed translational fusions of mEOS variants with five plasma membrane proteins, REM1.2, PLASMA MEMBRANE INTRINSIC PROTEIN 2;1 (PIP2;1), ARABIDOPSIS H^+^ ATPase 2 (AHA2), LOW TEMPERATURE INDUCED PROTEIN 6a (LTI6a) and the Rho of Plants 6 (ROP6) stably expressed in *Arabidopsis thaliana*. For all proteins, we observed that 488 nm illumination led to the recovery of fluorescent molecules (Supplementary Figure 2). Exposition to different light intensities indicated that approximately 10 µW is sufficient to recover half of the dark state mEOS molecules (Supplementary Figure 3). In transient assays we observed that upon 488 nm illumination, single molecules continuously emit light while they seemed to cycle between fluorescent and dark states in its absence (Supplementary Figure 1C, Supplementary Movie 1). Cyclic application of 488 nm light on seedlings expressing the relatively static PIP2;1-mEOS further suggests that the disappearance of fluorescent molecules is due to the re-entry into the long-lived dark state as molecules can be recovered at the same position during a second application of 488 nm light (Supplementary Figure 4). Altogether, these observations indicate that photo-converted mEOS molecules are predominantly in a non-fluorescent dark state and continuously cycle between a fluorescent and a long-lived dark state when expressed in plants. These observations also indicates that, due to the cycling between long-lived dark and fluorescent states, the same molecules are probably counted several times as independent molecules in classical PALM experiments. We observed that continuous 488 nm application maintained mEOS molecules in a fluorescent state (Supplementary Movie 1, Supplementary Movie 2) which we further quantitatively explored. By comparing the tracking duration of single molecules in presence and absence of 488 nm light exposure we observed that continuous illumination with 488 nm light increased the duration of single-molecule tracking (Supplementary Figure 5-7) without affecting the computed instantaneous diffusion coefficient (Supplementary Figure 8). Furthermore, we observed that continuous 488 nm illumination effectively prolonged tracking duration independently of the mEOS variants, the expression conditions, or the mobility behavior of the protein tested (Supplementary Figure 5-8). In these experiments we tracked molecules that were previously photo-converted and excited due to ambient and imaging conditions. Since fluorescent proteins emit a finite number of photons before photobleaching (Nguyen et al., 2023) we reasoned that we could track newly photo-converted molecules longer. Thus, we performed single-molecule tracking experiments upon low irradiation with 405 nm laser to induce mEOS photo-conversion, in absence and presence of 488 nm illumination. Under these imaging conditions (Figure 1C) we observed an additional increase in the proportion of long single-molecule tracks (longer than 5 seconds) representing approximately 15 % of all tracks (Supplementary Figure 9) upon 488 nm illumination. We then optimized the photon budget by reducing both the image acquisition frequency and the intensity of the 561 nm excitation laser. We imaged the cell-surface receptors BAK1 C-terminally fused to mEOS3.2 stably expressed in Arabidopsis. Under these conditions, we consistently observed and tracked individual receptors for durations that far exceed current gold-standard SMLM procedures in plants and animals (Figure 1 D-G). Together, these results demonstrate that tuning mEOS photophysics enables long-term single-molecule imaging and reliable tracking of individual molecules over seconds to minutes.

**Figure 1.**
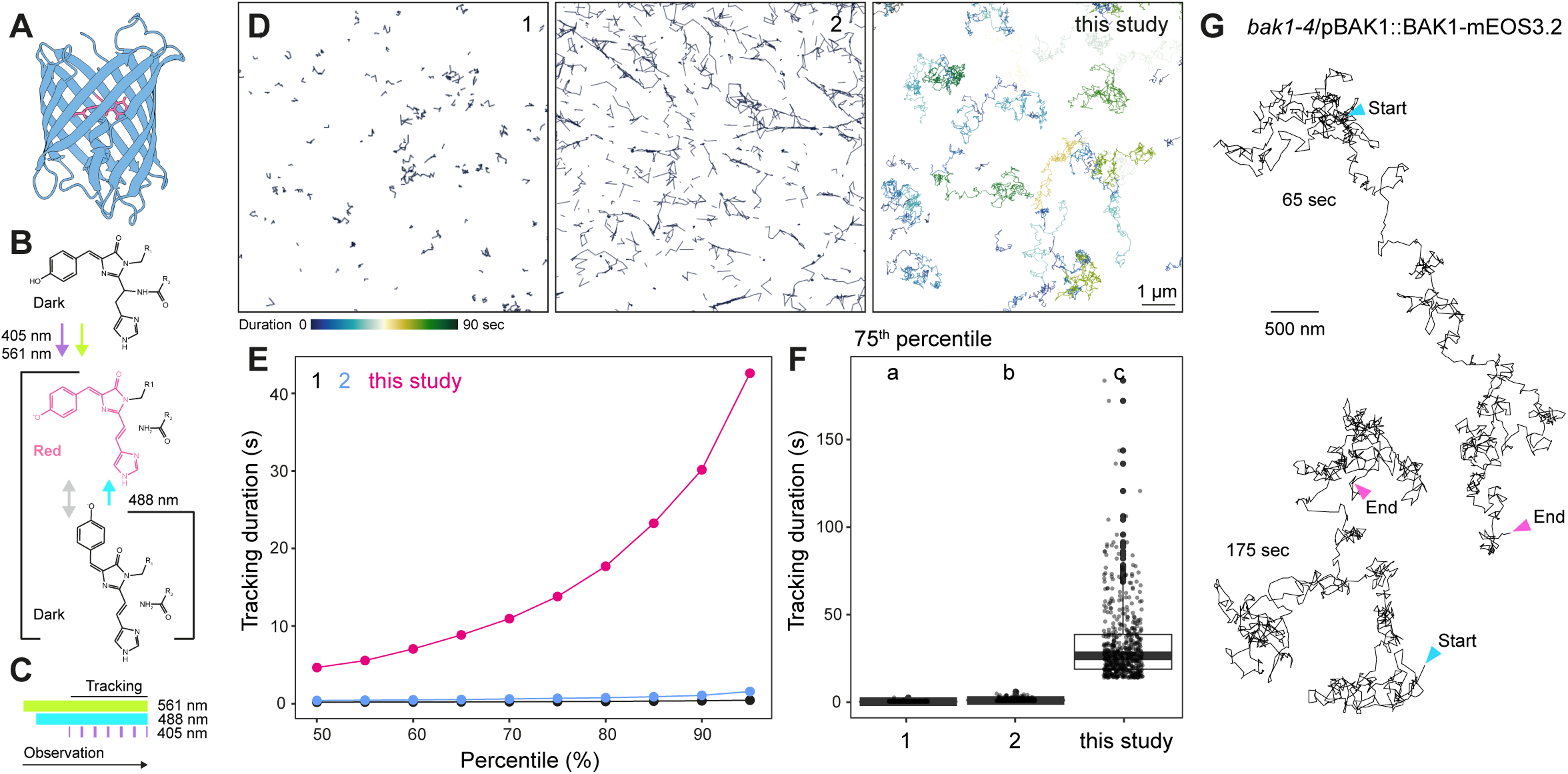
Minute-long tracking of single cell surface receptors. **A.** Structure (AlphaFold) of the fluorescent protein mEOS3.2 in which the beta barrel body is depicted in blue and the chromophore in pink. **B.** Selected photochromic transitions of mEOS chromophore. mEOS molecules are photoconverted using 405 nm light, excited using 561 nm light and spontaneously cycles between excited-fluorescent (red) and excited-dark state. They can be maintained in an excited-fluorescent state using 488 nm light. **C.** Illustration of the illumination conditions used for long-term single-molecule tracking. **D-F**. Representative images (**D**) and quantification (**E**-**F**) of single-molecules trajectories duration observed for Lti6a-mEOS2 in (Bayle et al., 2021)(1), MAP4- mEOS4b (De Zitter et al., 2019)(2) and BAK1-mEOS3.2 (this study). Single-molecule trajectories are colored based on their duration. Conditions that do not share a letter are significantly different in Wilcoxon rank sum test (p<0.005). **G**. Examples of very long-term single-molecule trajectories of BAK1-mEOS3.2.

Various imaging modalities have shown that the plasma membrane is laterally organized into nanodomains (Bücherl et al., 2017; Demir et al., 2013; Gronnier et al., 2022; Jarsch et al., 2014; Kleine-Vehn et al., 2011; Ma et al., 2022; Raffaele et al., 2009), which are proposed to provide means for the dynamic regulation of membrane-associated processes (Jaillais et al., 2024). However, the organization and dynamics of molecules are most often inferred from static images, diffraction-limited bulk fluorescence recovery measurements, or ensemble reconstructions in SMLM (Betzig et al., 2006; Shroff et al., 2007). The dynamic nature of the plant plasma membrane nano-organization has remained inaccessible. Our long-term single-molecule imaging modality revealed complex diffusion behavior that cannot adequately described by a single diffusion mode, as molecules frequently transitioned between phases of free diffusion and spatial arrest within the heterogenous plasma membrane (Figure 2) (Tabei et al., 2013; Weigel et al., 2011). Capturing such behavior requires computational tools that segment trajectories into distinct diffusive states, rather than assigning a single diffusion class to an entire track. (Magdziarz et al., 2009; Matsuda et al., 2018; Muñoz-Gil et al., 2021; Thapa et al., 2018; Vega et al., 2018).

**Figure 2.**
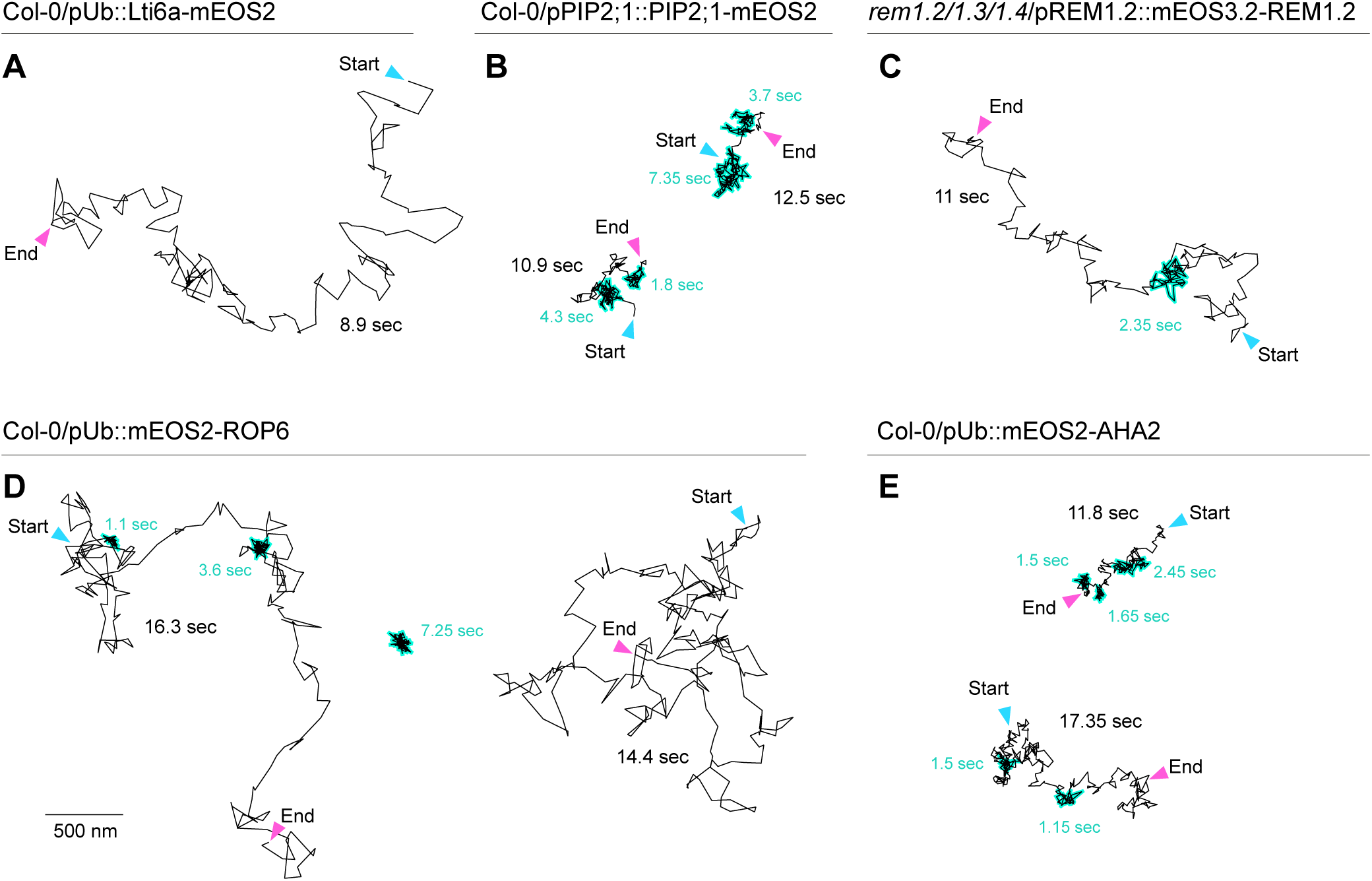
Live-cell nanoscale observations of single-molecule spatial arrests. Examples of single-molecule tracks obtained using photochromic recovery in stable *Arabidopsis* transgenic plants, Col-0/pUb10::Lti6a-mEOS2, Col-0/pUb::mEOS2-AHA2, Col-0/pPIP2;1::PIP2;1-mEOS2, Col-0/pUb::mEOS2-ROP6, and *rem1.2/rem1.3/rem1.4*/ pREM1.2::mEOS3.2-REM1.2. Imaging was performed in five-day-old seedlings cotyledon epidermal cells in presence of continuous 488 nm light exposure (25 µW). Apparent spatial arrests have been manually curated.

Diffusion analysis has been addressed using a spectrum of machine learning strategies, spanning classical methods such as hidden Markov models (HMMs) (Das et al., 2009; Matsuda et al., 2018; Ott et al., 2013; Persson et al., 2013) and random forests (RFs) (Muñoz-Gil et al., 2020) to modern deep learning–based approaches (Asghar et al., 2025; Dosset et al., 2016; Granik et al., 2019). While deep learning approaches often outperform classical machine learning methods, they have been reported to exhibit limited generalizability, frequently requiring extensive retraining and substantial computational resources. These constraints can hinder their broader adoption by non-expert users and can result in longer processing times (Kowalek et al., 2019). To robustly detect and analyse spatial arrest events within single molecule trajectories, we developed Computational Analysis of Spatial Arrest (CASTA), a hybrid machine learning- and rule-based decision framework (Figure 3 A-C). Available as an easy to setup and use Python package, CASTA enables high-throughput analysis of large single-particle tracking data sets. Because CASTA was designed for binary classification of diffusive versus spatially arrested track segments, it does not require *a priori* assumptions about the number or nature of underlying diffusion states, nor does it impose constraints in terms of minimum number of trajectories or maximum number of timepoints for reliable classification as required for other methods (Asghar et al., 2025; Persson et al., 2013). Moreover, CASTA can detect changes between states with as little as 10 timepoints, which is lower compared to other methods (Dosset et al., 2016; Vega et al., 2018). At the core of the pipeline is a Hidden Markov Model (HMM) (Rabiner, 2002) which probabilistically classifies each trajectory segment as either exhibiting confined or free diffusion (Figure 3B, Supplementary Figure 10A, B). To train the HMM model, we generated ground-truth data by simulating single-particle trajectories using the Anomalous Diffusion Challenge (AnDi) framework (Muñoz-Gil et al., 2021) (Figure 3F). Simulation parameters, such as diffusion rates and the distribution of spatial arrest area were chosen to recapitulate the broad range of experimentally observed plant plasma membrane protein dynamics (Figure 3D, E). In addition, to complement the HMM and enhance classification robustness, we integrated quantitative features, termed diffusional signature metrics, that capture distinct local properties of single particle motion (Figure 3B, Supplementary Figure 10C, D). For each time point, the HMM prediction and the four metric-based classifications are combined in a majority-voting scheme (Figure 3B). If at least three of the five criteria agree, the corresponding state (confined or diffusive) is assigned. Spatial arrests are subsequently predicted based on a minimum of 5 consecutive steps classified as confined. Finally, a convex hull is constructed around all consecutive confined steps to define the area of the arrest event (Figure 3C). To benchmark its performance, we tested CASTA on 480 distinct simulations each having a unique combination of diffusional parameters, spanning a broad spectrum of diffusional behaviors (Supplementary Table 4). We found that CASTA captured spatial arrests with an average precision exceeding 97 % within the parameter range typically observed for plasma membrane-localized proteins (Figure 3G, I). Moreover, CASTA robustly detected changes in diffusional behavior with as little as 5-fold difference between freely diffusing and confined states (Figure 3G), whereas previously reported values typically range from 10- to 100-fold (Daumas et al., 2003). CASTA also reliably detected spatial arrest with as few as 12 localizations (Supplementary Figure 11A), below the minimum required to detect state changes in otherexisting approaches (Dosset et al., 2016; Vega et al., 2018). Due to the inherent noise of single-particle tracking data, CASTA was conservatively tuned to prioritize high precision, at the moderate expense of reduced recall (∼ 70 %) for parameter regimes that deviate from in vivo single-molecule behavior (e.g., large anomalous exponents and confined areas) (Figure 3 H, J, Supplementary Figure 11B). Finally, we compared CASTA with DC-MSS (Vega et al., 2018), which has previously been used for classification of diffusion types of plant PM proteins(Rohr, Rausch, et al., 2024). We found that CASTA outperformed DC-MSS in terms of precision in a broad range of scenarios (Supplementary Figure 11B). Together, these results establish CASTA as a sensitive and precise analysis pipeline for detecting spatial confinement events, enabling robust quantitative analysis of nanoscale membrane dynamics.

**Figure 3.**
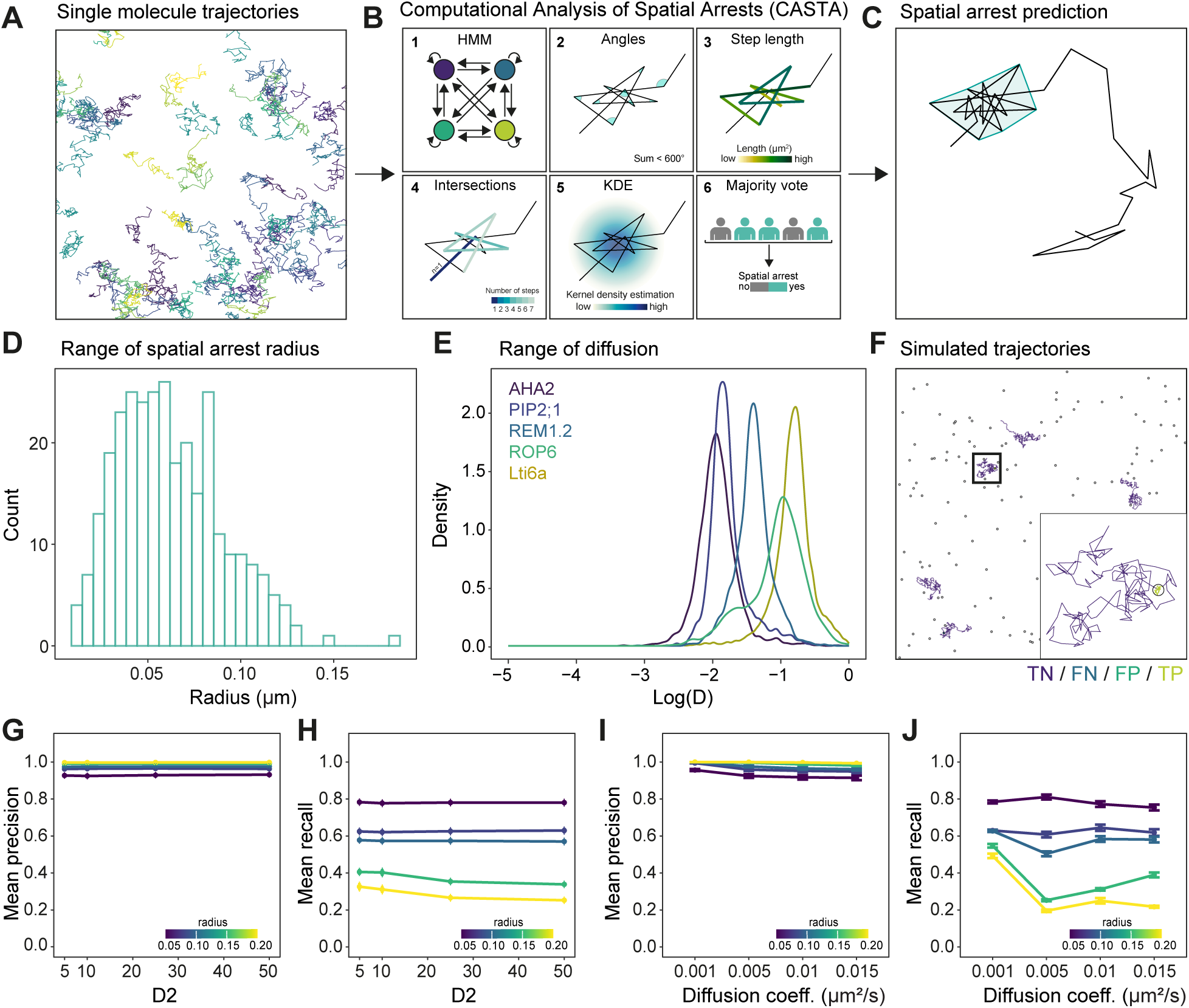
Computational analysis of spatial arrest (CASTA). **A-C.** Illustration of CASTA. Single-molecule trajectories (**A**) are subjected to a hybrid machine learning- and rule-based decision pipeline that consists of trained Hidden Markov Model (HMM, 1) and four diffusional signature metrics (2-5) which are combined in a majority-voting scheme (6) to predict spatial arrest (**C**). **D.** Distribution of the area of spatial arrest events measured from single-molecule trajectories. **E.** Analysis of the of the apparent diffusion coefficient D of mEOS2-AHA2, PIP2;1-mEOS2, mEOS3.2-REM1.2, mEOS2-ROP6 and Lti6a-mEOS2 molecules, observed in five-days-old Arabidopsis epidermal cells. Total number of cells (n) analysed equal 12 for mEOS2-AHA2, n=11 PIP2;1-mEOS2, n=19 for mEOS3.2-REM1.2, n=9 for mEOS2-ROP6 and n=11 for Lti6a-mEOS2, from two independent experiments. **F.** Analysis by CASTA of ground truth simulated single-particle trajectories. The zoomed-in region depicts a single trajectory entering and leaving a confinement zone (black circle). Trajectory segments are color-coded based on prediction performance where (TN) = true negatives, (FN) = false negatives, (FP) = false positives and (TP) = true positives. **G-J.** Mean precision (**G** and **I**) and recall (**H** and **J**) performance of CASTA on the confined class in benchmarking simulations depending on the fold of diffusion rate (D2) between inside and outside compartments (**G-H**) and diffusion coefficient (**I-J**) for five distinct compartment radii.

Finally, we used CASTA to analyse the behavior of a representative set of plasma membrane proteins. We detected spatial arrest events for all five tested proteins (Figure 4B). This contrasts with the bulk analysis of the diffusion coefficient, as usually performed in spt-PALM experiments, which failed to reveal the distinctive mobility behavior except for ROP6 (Fig. 3E). We computed the frequency of spatial arrest events, their duration and the corresponding area for each protein (Figure 4C). These analyses revealed the distinctive diffusional behavior of plasma membrane proteins. For instance, while spatial arrests of Lti6a were rare, they tended to last longer than those of the other tested proteins. Combining Photochromic reversion and CASTA, we recently unveil the nanoscale spatial and temporal logic underlying the formation of leucine-rich repeat receptor kinase complexes (von Arx et al., 2026).

**Figure 4.**
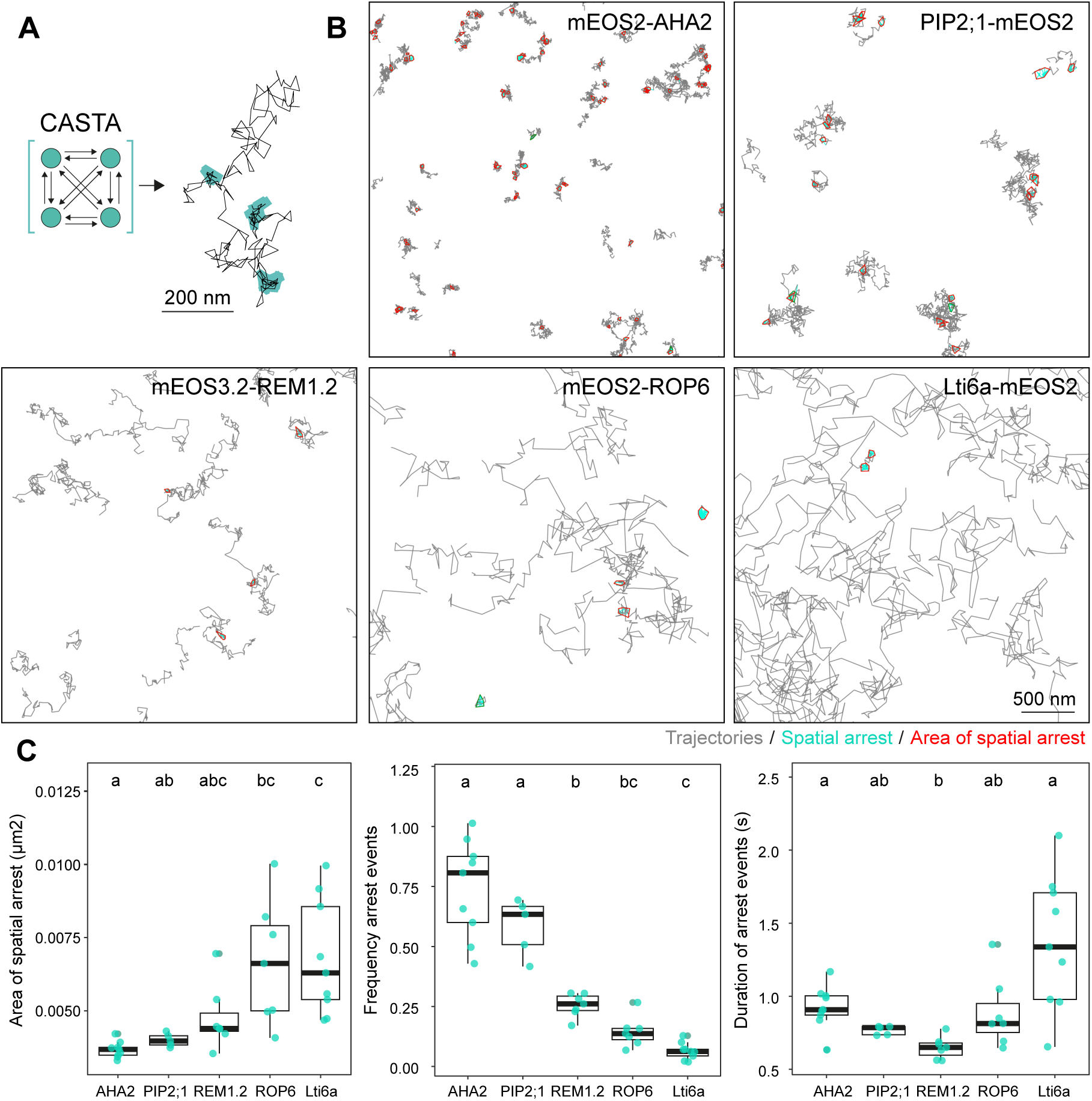
CASTA highlights distinctive spatial arrest behavior of plasma membrane proteins. **A**. Schematic depiction of the analysis of spatial arrest events by CASTA within single-molecule tracks. **B**. Examples of trajectory segmentations using CASTA on long-term single-molecule tracks observed in the stable *Arabidopsis* transgenic plants Col-0/pUb10::Lti6a-mEOS2, Col-0/pUb::mEOS2-AHA2, Col-0/pPIP2;1::PIP2;1-mEOS2, Col-0/pUb::mEOS2-ROP6, and *rem1.2/rem1.3/rem1.4*/pREM1.2::mEOS3.2-REM1.2. Imaging was performed in five-day-old seedlings cotyledon epidermal cells in in presence of continuous 488 nm (25 µW) and continuous 561 nm (1000 µW), and in presence of intermittent 405 nm (2 µW) light exposure. Tracks segments predicted as transient STA are indicated in red. **C**. Quantitative analysis of the frequency of spatial arrest events per trajectory (left) their duration (middle) and the area in which molecules were spatially arrested. Graphs are boxplots, scattered data points indicate average values obtained from individual cells. Total number of cells (n) analysed equal 9 for Lti6a-mEOS2, n=10 for mEOS2-AHA2, n=7 for mEOS2-ROP6, n=5 for PIP2;1-mEOS2 and n=7 for mEOS3.2-REM1.2. Conditions which do not share a letter are significantly different in Wilcoxon rank sum test (p<0.005).

In sum, by exploiting photophysical properties of mEOS variants, we overcame a long-standing limitation of single-molecule imaging, allowing for long-term single-molecule imaging and the direct observation of dynamic molecular events at the plasma membrane. We provide a computational solution for the automated analysis of single-molecule nanoscale dynamics. This integrated experimental and computational framework thus opens new avenues for mechanistic studies of membrane organization and signaling dynamics.

## Acknowledgement

We thank all members of the NanoSignaling Laboratory for fruitful discussions and comments on the manuscript. We thank Alexandre Martinière for providing the Col-0/pUb::Lti6a-mEOS2, Col-0/pPIP2;1::PIP2;1-mEOS2, Col-0/pUb::mEOS2-ROP6 and Col-0/pUb::mEOS2-AHA2 transgenic lines. We thank Sébastien Mongrand for providing the rem1.2/rem1.3/rem1.4 mutant. We thank Vincent Bayle, Alexandre Martinière, Dominique Bourgeois and Yvon Jaillais for helpful discussions and comments on the manuscript. This research was supported by the Deutsche Forschungsgemeinschaft (DFG) grants SFB1101-A09 and TRR356-B01 to J.G and Z02-SFB1101 to S.z.O.K.

---

**Supplementary Figure 1.**
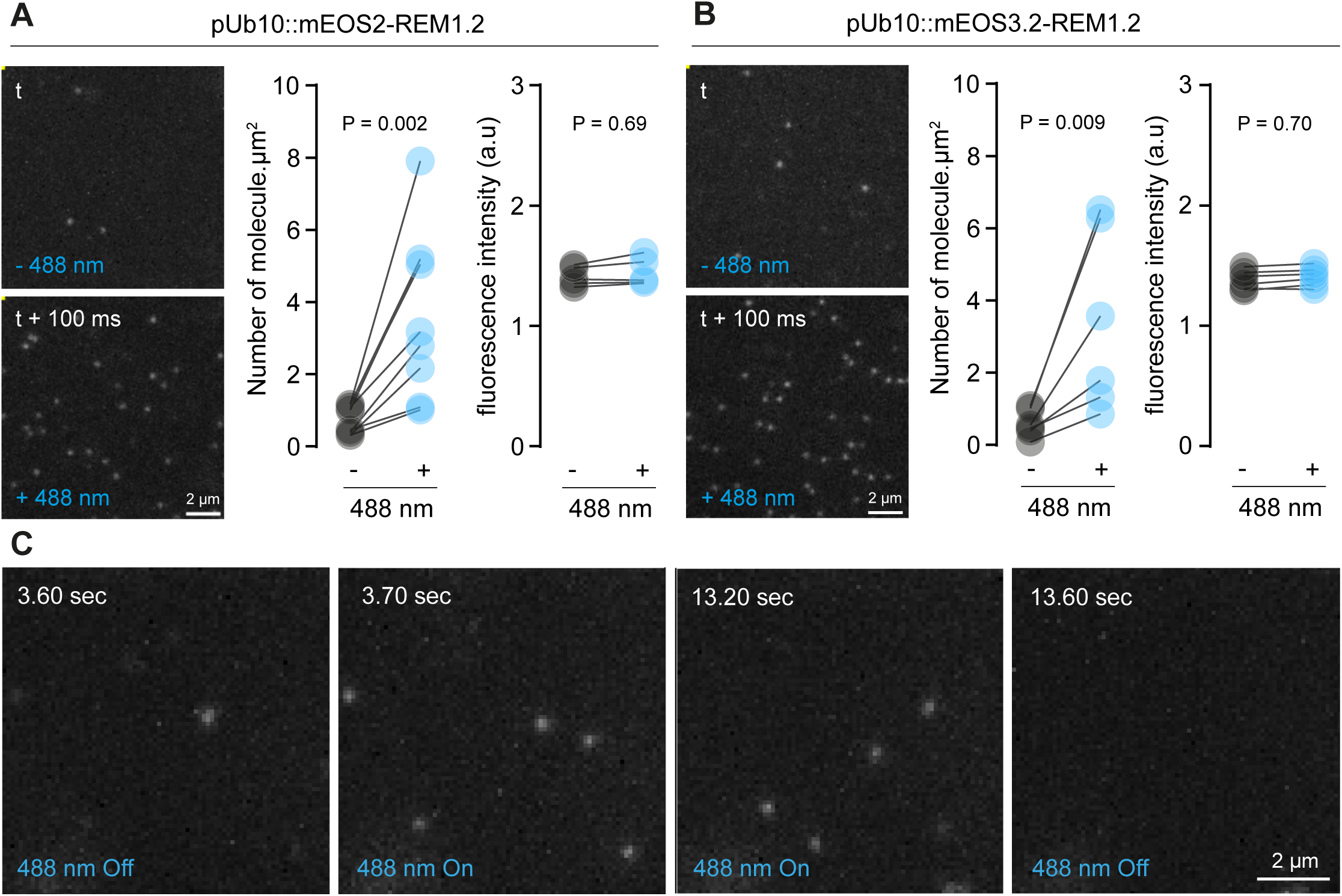
Effect of 488nm illumination on mEOS2-REM1.2 and mEOS3.2-REM1.2 molecules transiently expressed in *N. benthamiana*. Quantification of the number of molecules observed and of their fluorescence intensity before (t) and upon 488 nm light exposure (t + 100ms) for mEOS-REM1.2 (**A**) and mEOS3.2-REM1.2 (**B**) molecules transiently expressed in *N. benthamiana* 30 hours post infiltration. Each dot represents the average value observed for individual cells. The lines connect values observed for the same cell before and upon 488 nm laser application. P values report two-tailed Mann-Whitney statistical test. **C**. Images correspond to individual time frames of mEOS3.2-REM1.2 molecules observed upon transient expression in *N. benthamiana*. See Supplementary Movie 1.

**Supplementary Figure 2.**
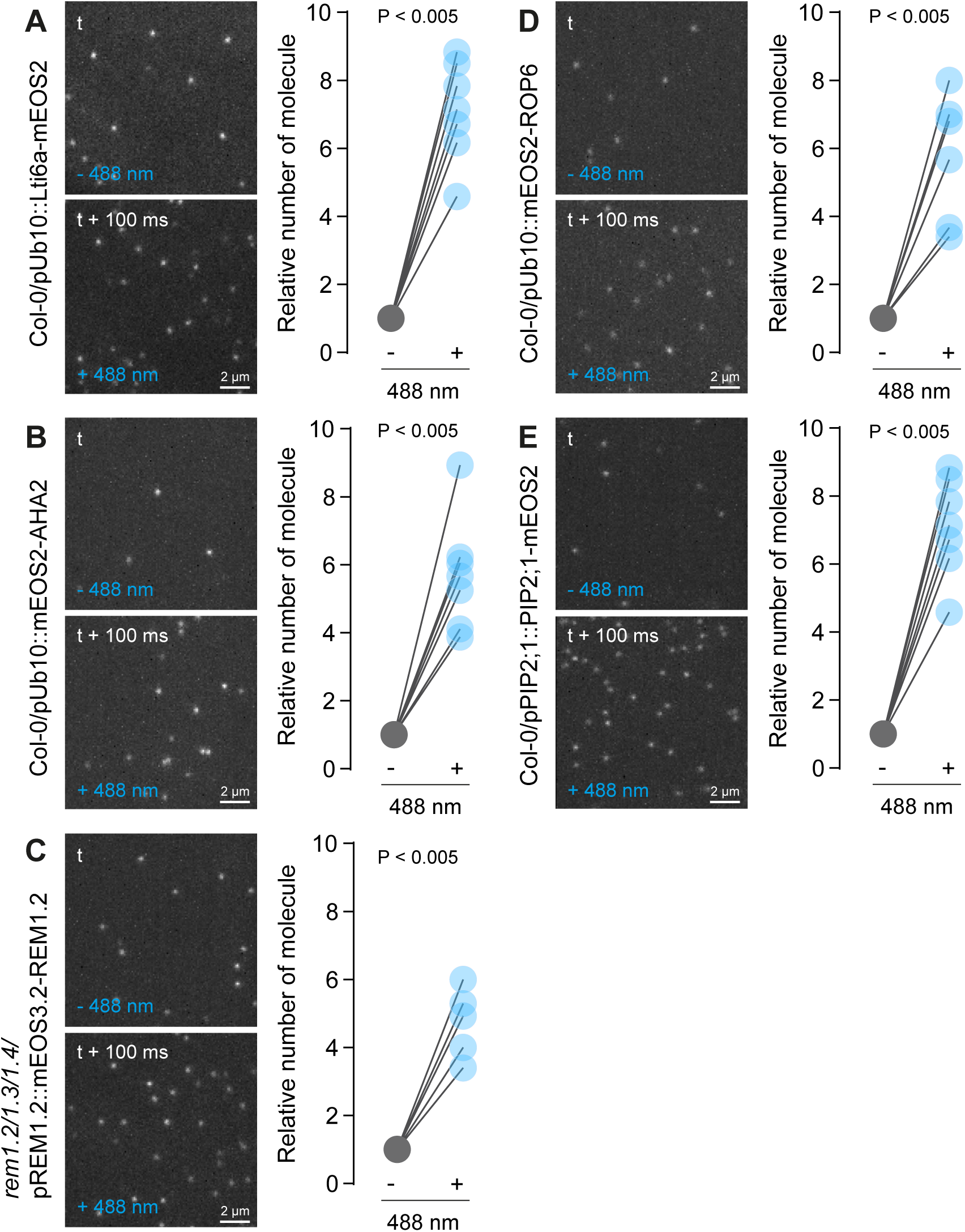
Effect of 488nm illumination on five plasma membrane-localized mEOS translational fusions expressed in *Arabidopsis thaliana*. Quantification of the number of molecules observed before (t) and upon 488 nm light exposure (t + 100ms) for Lti6a-mEOS2 (**A**) mEOS2-AHA2 (**B**), mEOS3.2-REM1.2 (**C**), mEOS2-ROP6 (**D**), and PIP2;1-mEOS2 (**E**) in five-days-old Arabidopsis cotyledon epidermal cells. Each dot represents the average value observed for individual cells expressed as relative to the control condition (- 488 nm light application). The lines connect values observed for the same cell before and after 488 nm laser application. P values report two-tailed Mann-Whitney statistical test.

**Supplementary Figure 3.**
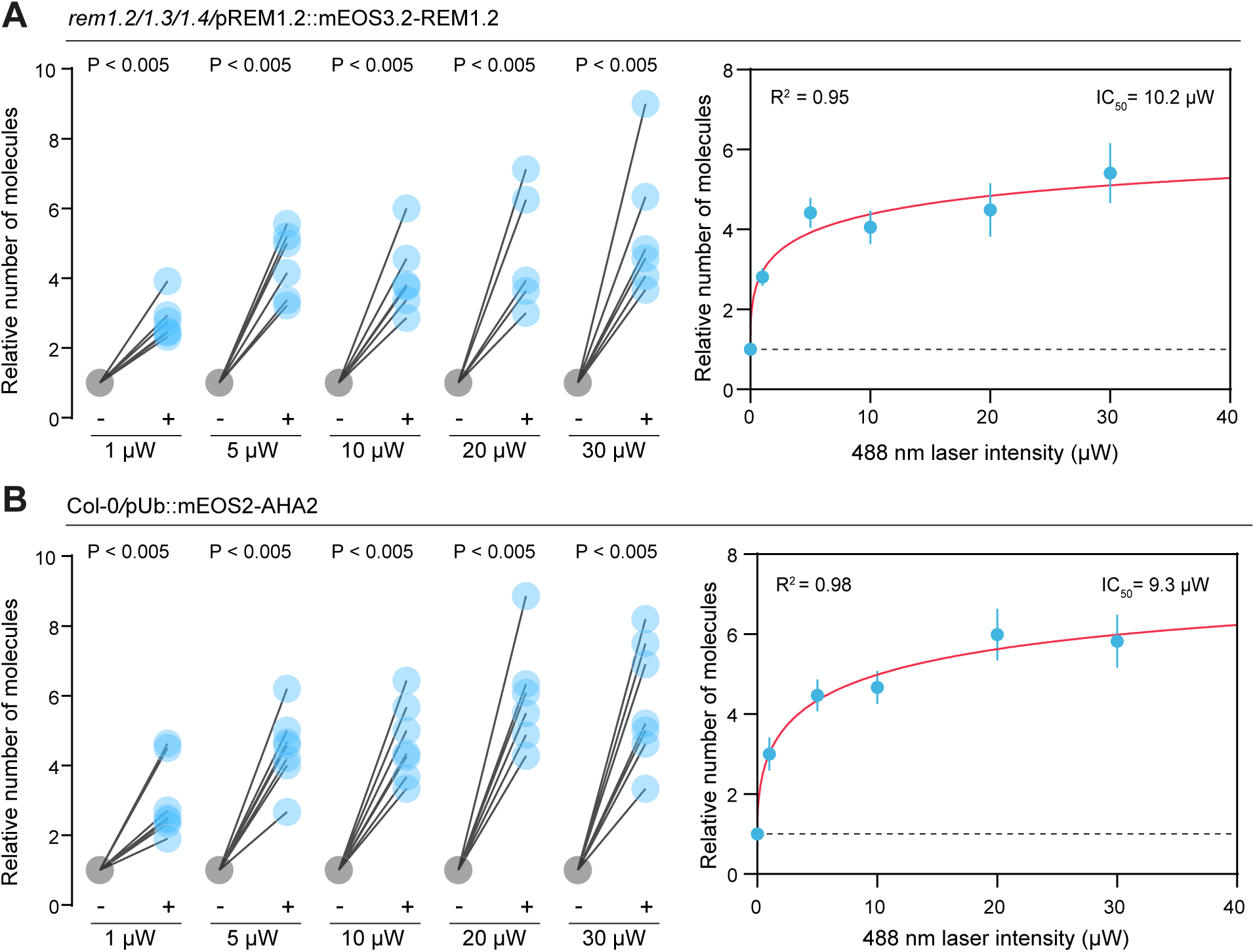
Analysis of the dose response to 488 nm light exposure. Quantification of the number of mEOS3.2-REM1.2 (**A-B**) and mEOS2-AHA2 (**C-D**) molecules observed before and upon exposure to 488 nm light of several intensities in five-days-old Arabidopsis epidermal cells. In **A** and **C**, each dot represents the average value observed for individual cells expressed as relative to the control condition (without 488 nm light application). The lines connect values observed for the same cell before and upon 488 nm laser application. P values report two-tailed Mann-Whitney statistical test. The graphs in (**B**) and (**D**) depict the number of observed molecules as a function of 488 nm light intensity. Data points represent average values observed across cells for each light intensity conditions ± standard error to the mean. Red curves represent non-linear fits of the data. Total number of cells (n) analysed equal 56 for mEOS3.2-REM1.2 and n=57 for mEOS2-AHA2.

**Supplementary Figure 4.**
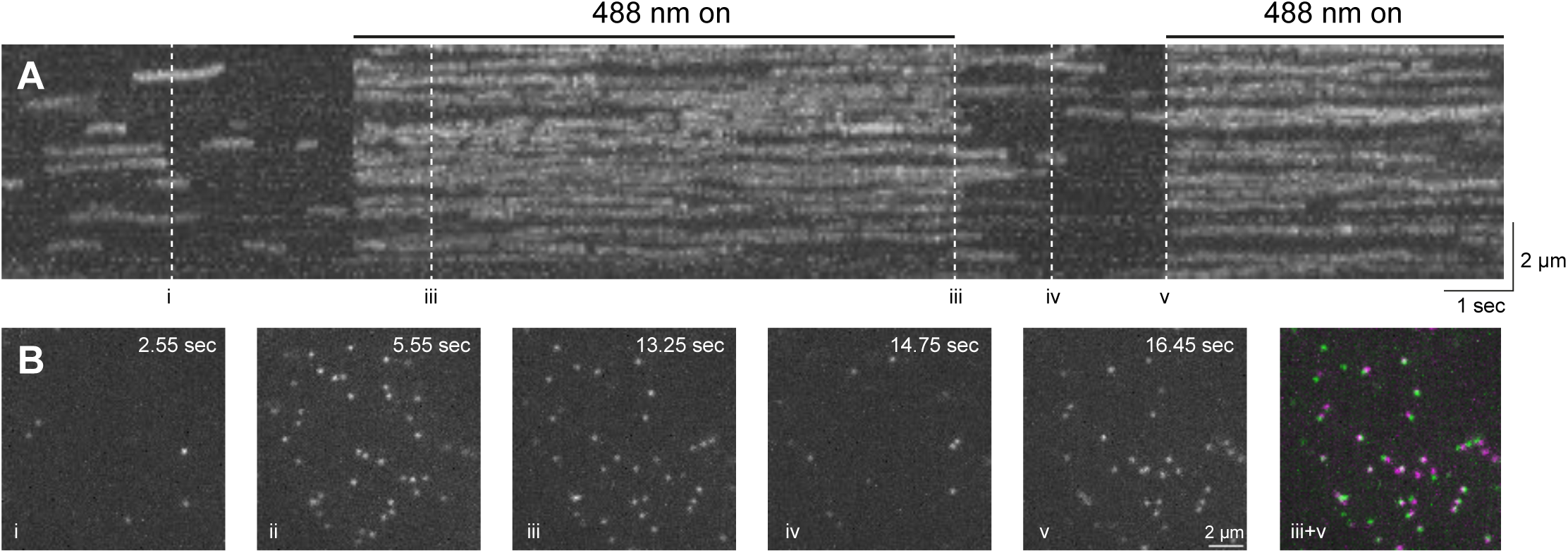
Sequential exposure to 488 nm shows controlled cycling of photo-converted mEOS molecules between fluorescent and dark state. Analysis of PIP2;1-mEOS2 fluorescence overtime in five-days-old Arabidopsis epidermal cells. **A.** The image corresponds to a kymograph in which the fluorescence observed across x and y coordinates is projected a single plane (coordinate x) overtime. **B.** Each image corresponds to individual time frames (x,y) of the temporal projection depicted in A.

**Supplementary Figure 5.**
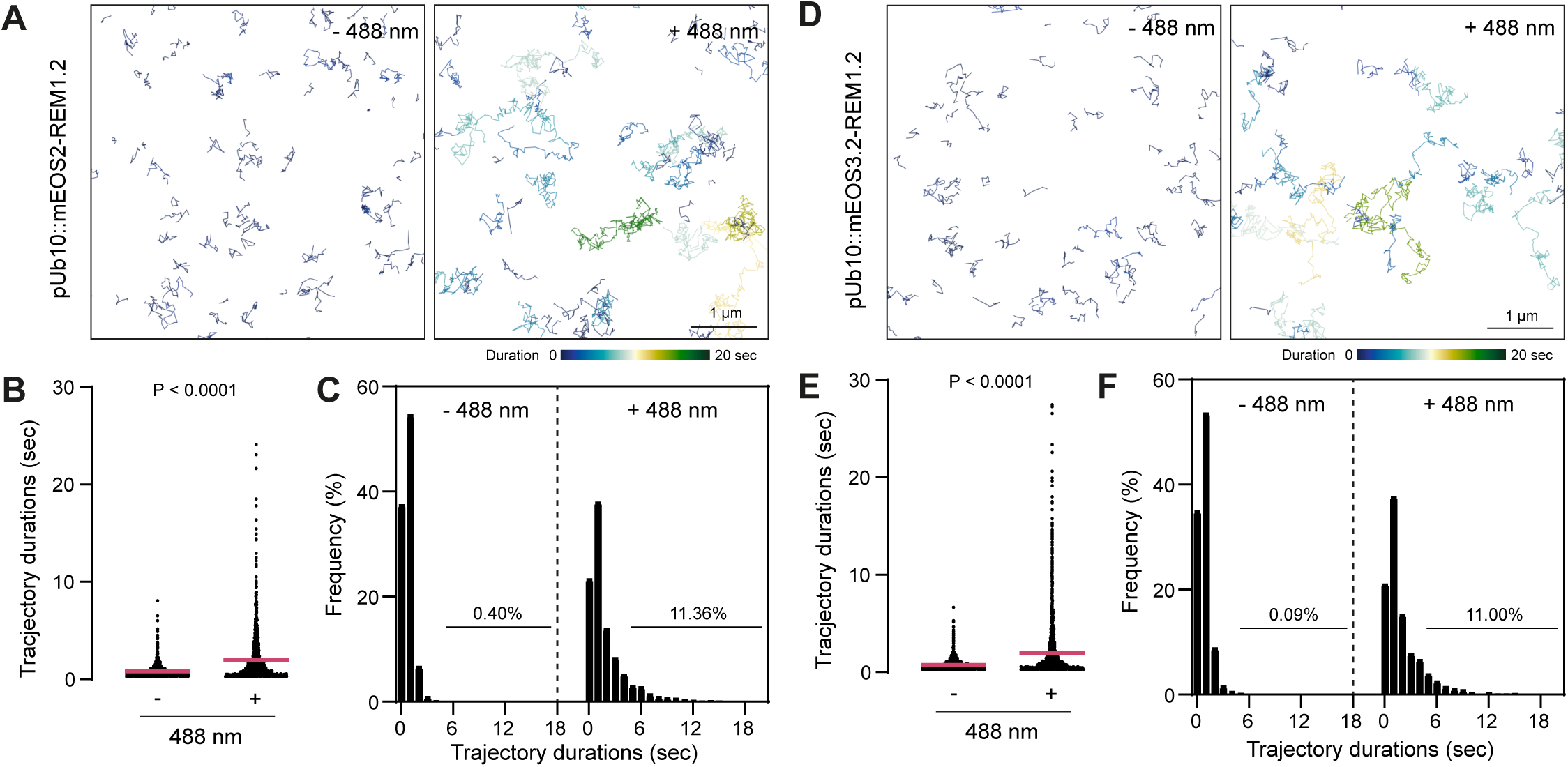
Effect of 488 nm light exposure on trajectory durations of mEOS2-REM1.2 and mEOS3.2-REM1.2 transiently expressed in *N. benthamiana*. **A** and **D** show representative images of single-molecule trajectories of mEOS2-REM1.2 (**A**) and mEOS3.2 (**D**) observed in 3-week-old *N. benthamiana* leaf epidermal cells in presence and absence of continuous 488 nm light exposure (25 µW). Trajectories are coloured based on their duration. **B** and **E**. Graphs depict trajectory durations for individual tracks of mEOS2-REM1.2 (**B**) and mEOS3.2- REM1.2 (**E**) in presence and absence of continuous 488 nm light exposure (25 µW). P values report two-tailed Mann-Whitney statistical test. **C** and **F**. Graphs correspond to normalized histograms of trajectory durations observed in presence and absence of continuous 488 nm light exposure (25 µW) mEOS2-REM1.2 (**C**) and mEOS3.2-REM1.2 (**F**). Percentages indicate the proportion of trajectories whose duration is equal or longer than 5 seconds. Total number of cells (n) analysed equal 25 for mEOS2-REM1.2, n=32 for mEOS3.2-REM1.2 from two independent experiments.

**Supplementary Figure 6.**
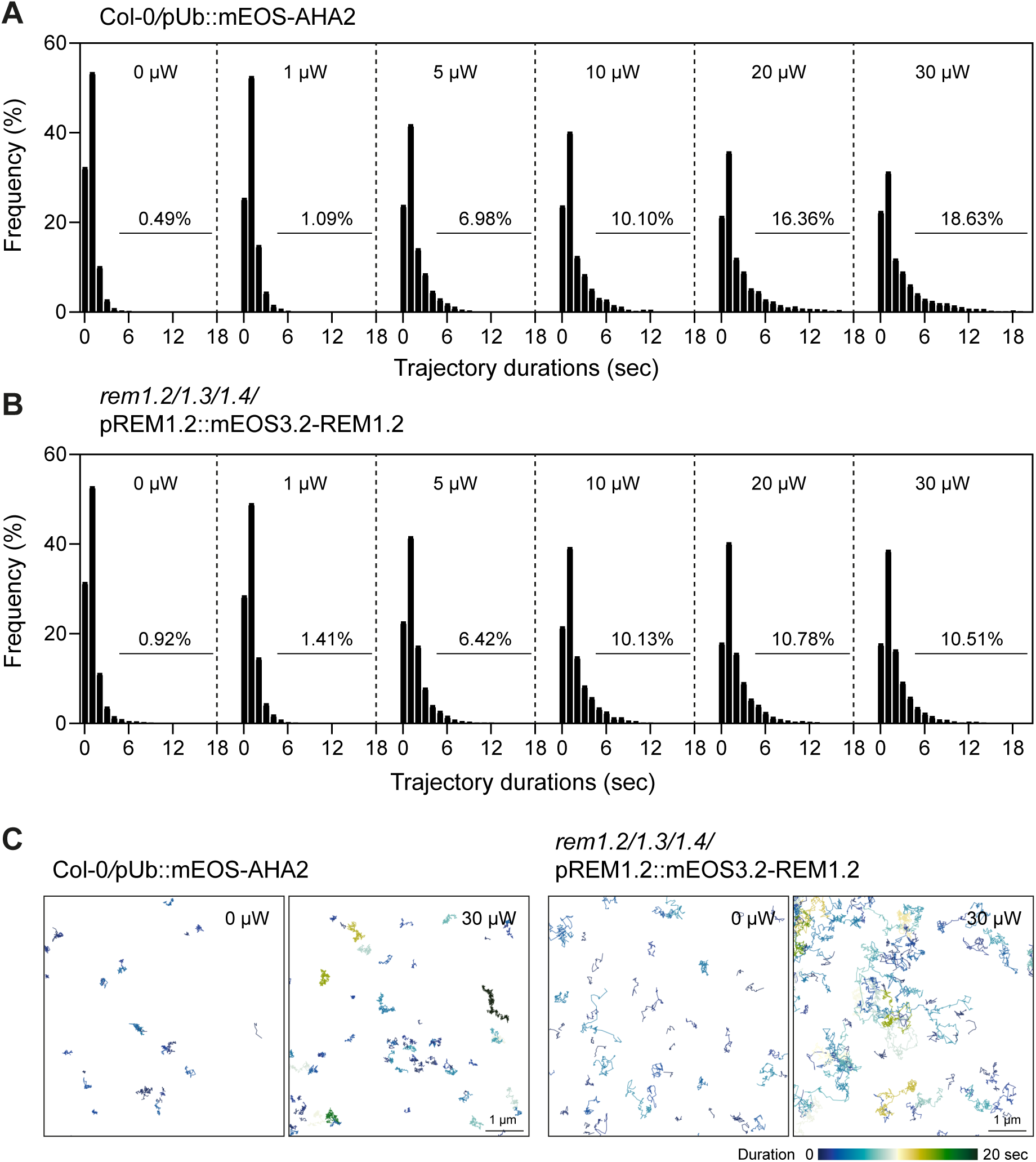
Effect of 488 nm light intensity on trajectory durations of mEOS2-AHA2 and mEOS3.2-REM1.2. **A-B**. Graphs correspond to normalized histograms of trajectory durations (s) observed upon different intensity of 488 nm light exposure for mEOS2-AHA2 (**A**) and mEOS3.2-REM1.2 (**B**). Percentages indicate the proportion of trajectories whose duration is equal or longer than 5 seconds. Total number of cells (n) analysed equal 35 for mEOS3.2-REM1.2 and n=38 for mEOS2-AHA2 collected in one experiment. **C.** Representative images of single-molecule trajectories of mEOS2-AHA2 (left) and mEOS3.2-REM1.2 (right). Trajectories are coloured based on their duration.

**Supplementary Figure 7.**
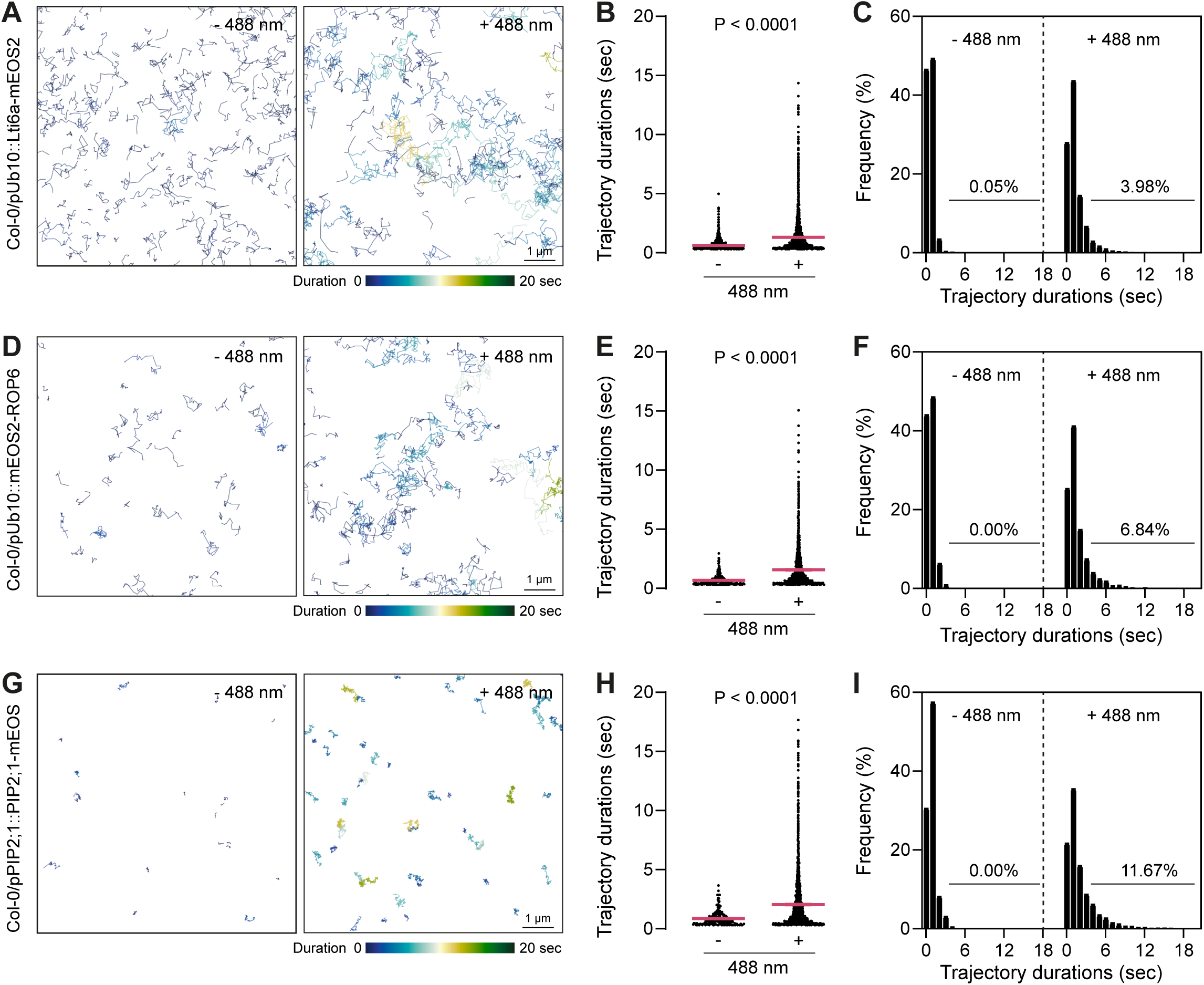
Effect of 488 nm light exposure on trajectory durations of Lti6a-mEOS2, mEOS2-ROP6 and PIP2;1-mEOS2. **A**, **D** and **G** show representative images of single-molecule trajectories of Lti6a-mEOS2 (**A**), mEOS2-ROP6 (**D**) and PIP2;1-mEOS2 (**G**) observed in five-days-old Arabidopsis epidermal cells in presence and absence of continuous 488 nm light exposure (25 µW). Trajectories are coloured based on their duration. **B**, **E** and **H**. Graphs depict trajectory durations for individual tracks of Lti6a-mEOS2 (**B**), mEOS2-ROP6 (**E**) and PIP2;1-mEOS2 (**H**) in presence and absence of continuous 488 nm light exposure (25 µW). P values report two-tailed Mann-Whitney statistical test. **C**, **F** and **I**. Graphs correspond to normalized histograms of trajectory durations observed in presence and absence of continuous 488 nm light exposure (25 µW) for Lti6a-mEOS2 (**C**), mEOS2-ROP6 (**F**) and PIP2;1-mEOS2 (**I**). Percentages indicate the proportion of trajectories which duration is equal or longer than 5 seconds. Total number of cells (n) analysed equal 18 for Lti6a-mEOS2, n=12 for mEOS2-ROP6 and n=11 for PIP2;1-mEOS2 from two independent experiments.

**Supplementary Figure 8.**
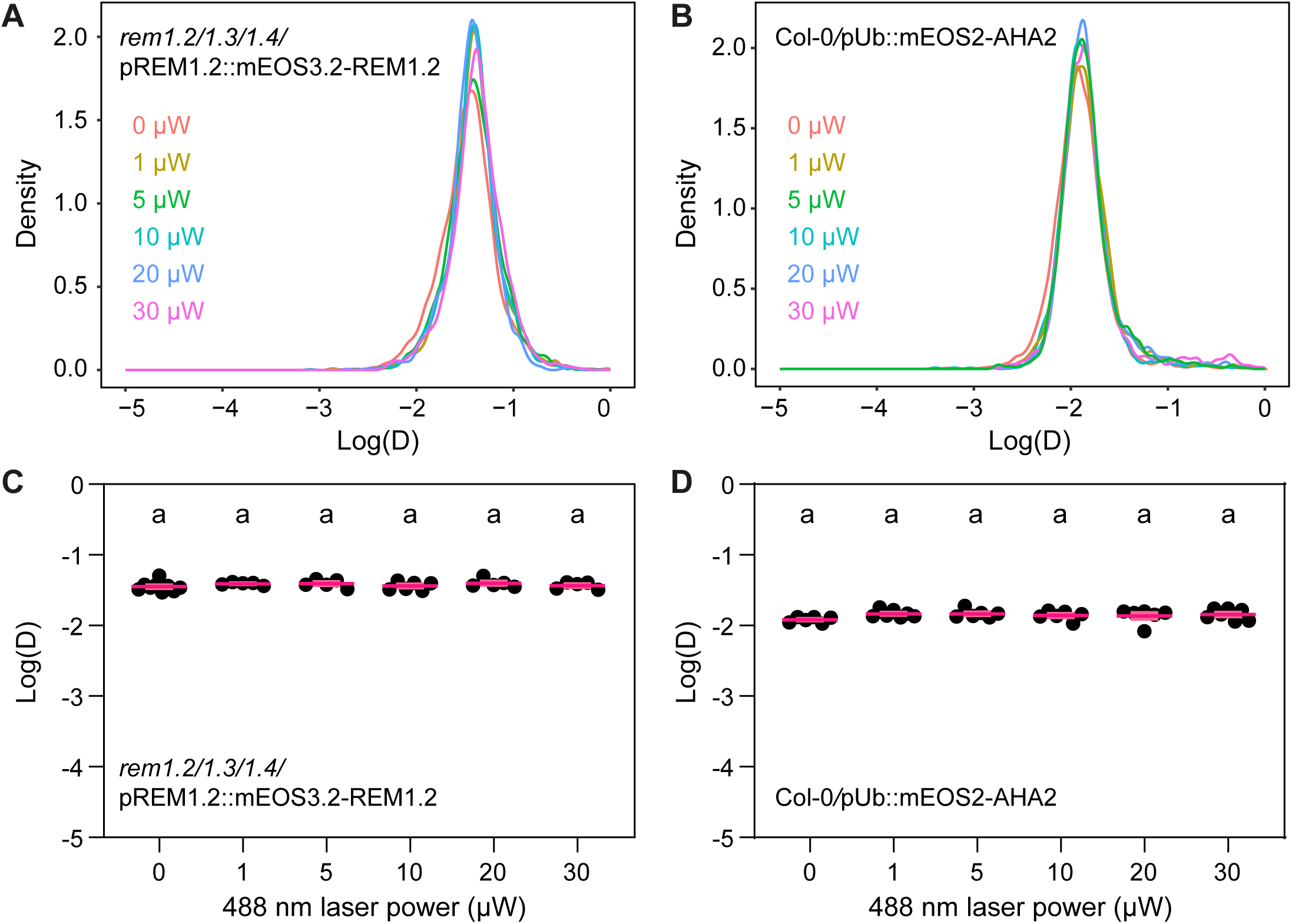
488 nm light exposure does not affect diffusion coefficient. Analysis of the diffusion coefficient (D) of mEOS3.2-REM1.2 (**A-C**) and mEOS2-AHA2 (**B-C**) upon exposure of five-days-old Arabidopsis epidermal cells with different 488 nm light intensities. **A** and **B** depict the distribution of mEOS3.2-REM1.2 and mEOS2-AHA2 molecules according to their apparent diffusion coefficient. **c** and **d** represent the average diffusion coefficient for individual cells. Total number of cells (n) analysed equal 35 for mEOS3.2-REM1.2 and n=38 for mEOS2-AHA2 collected in one experiment, pink crosses indicate mean ± s.e.m. Conditions sharing a letter are not significantly different in Dunn’s multiple comparison test (P>0.22).

**Supplementary Figure 9.**
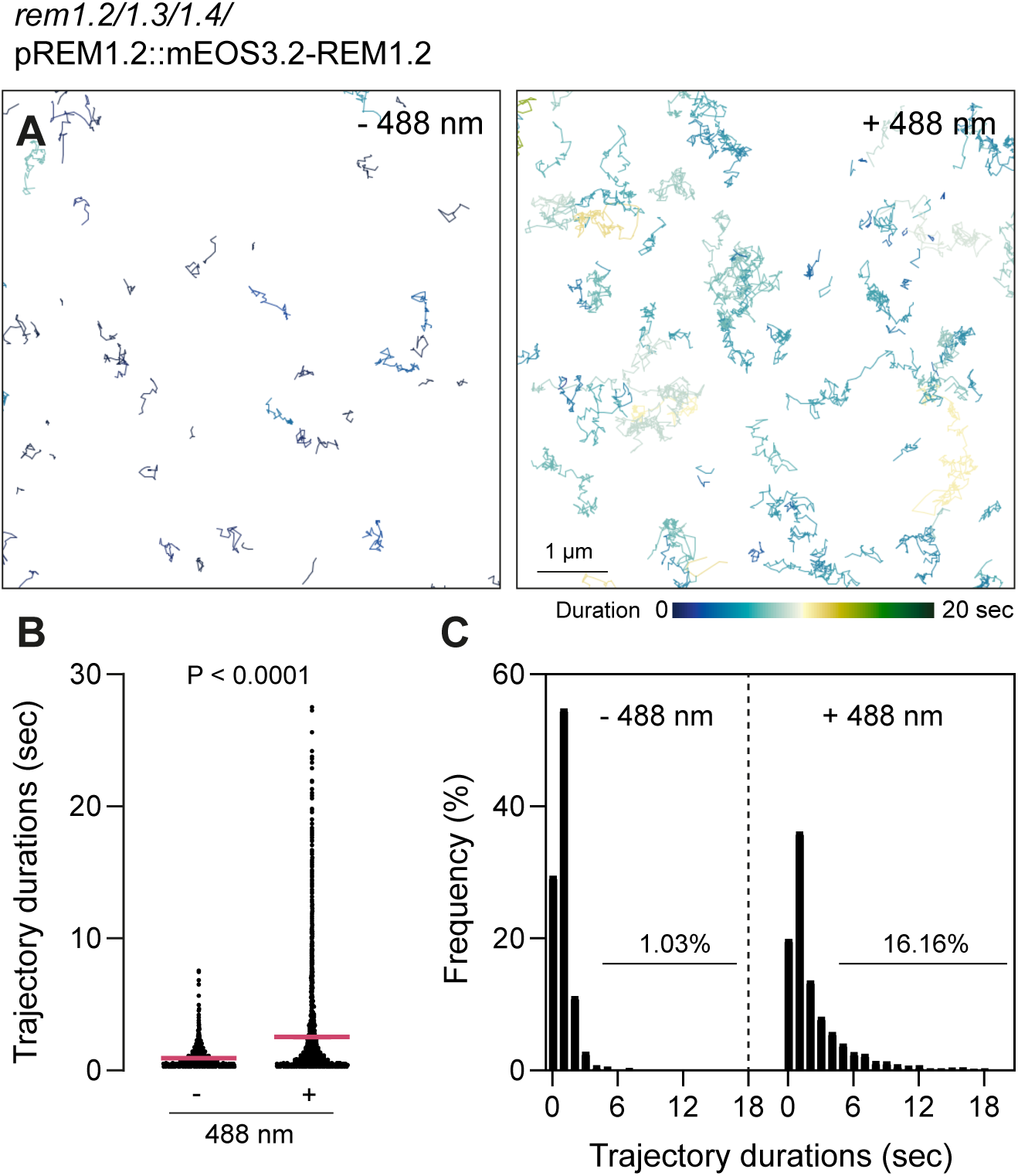
Analysis of trajectory duration of newly photo-converted molecules. **A.** Representative images of single molecule trajectories of mEOS3.2-REM1.2 observed in five-days-old Arabidopsis epidermal cells in presence and absence of continuous 488 nm light exposure (25 µW). **B.** Graph depicts trajectories duration for individual tracks mEOS3.2-REM1.2 in presence and absence of continuous 488 nm light exposure (25 µW). P values report two-tailed Mann-Whitney statistical test. **C.** Graphs correspond to normalized histogram of trajectories duration observed in presence and absence of continuous 488 nm light exposure (25 µW) for mEOS3.2-REM1.2. Percentages indicate the proportion of trajectories whose duration is equal or longer than 5 seconds. Total number of cells (n) analysed equal 18 from two independent experiments.

**Supplementary Figure 10.**
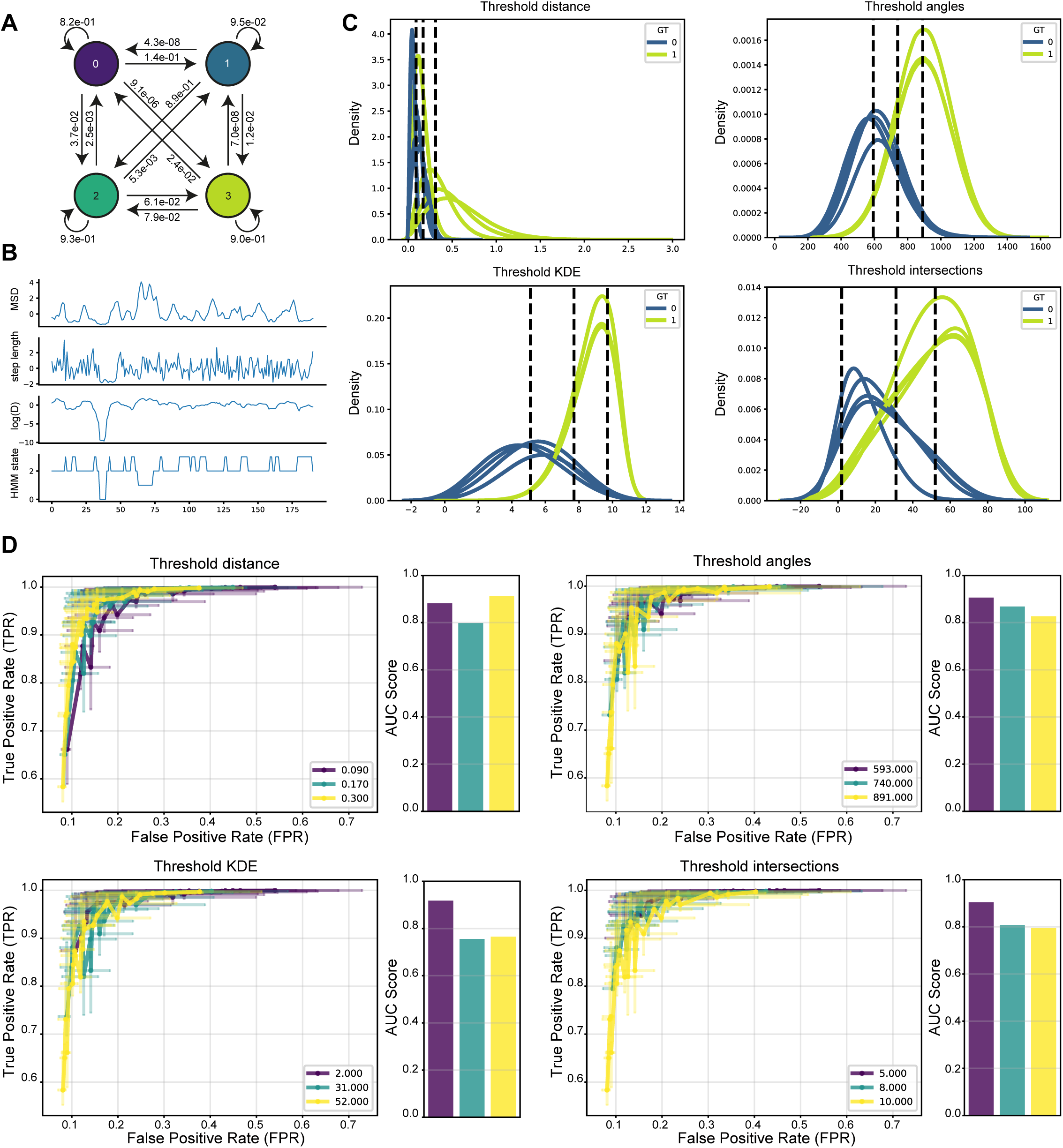
Hidden Markov Model and simulation-based evaluation of diffusional signature metrics for spatial arrest detection. **A.** Schematic depiction of Hidden Markov Model (HMM) hidden states and associated transition probabilities between states. The zero state is counted as the confined state, while one, two and three are not confined. **B.** HMM input features and hidden state output of an exemplary track. **C.** Density plots showing the distribution of diffusional feature values measured at each step in a track across multiple simulations (each N = 23 500 trajectories) with varying simulation parameters, grouped by the ground-truth value of the step (0 = confined, 1 = not confined). Dashed lines indicate candidate threshold values for the diffusional feature, the lowest and highest being the mean value of the feature values across the positive and negative group, and the middle being the value that optimally splits the two distributions according to Youden’s J statistic. **D.** True Positive Rate (TPR) and False Positive Rate (FPR) values for the same simulations as **B,** grouped for each feature and candidate threshold value, with error bars showing the distribution of the 25^th^ – 75^th^ quantile of values in both dimensions. Bar plots show the corresponding Area Under the ROC (Receiver Operating Characteristic) curve (AUC) score for each threshold value.

**Supplementary Figure 11.**
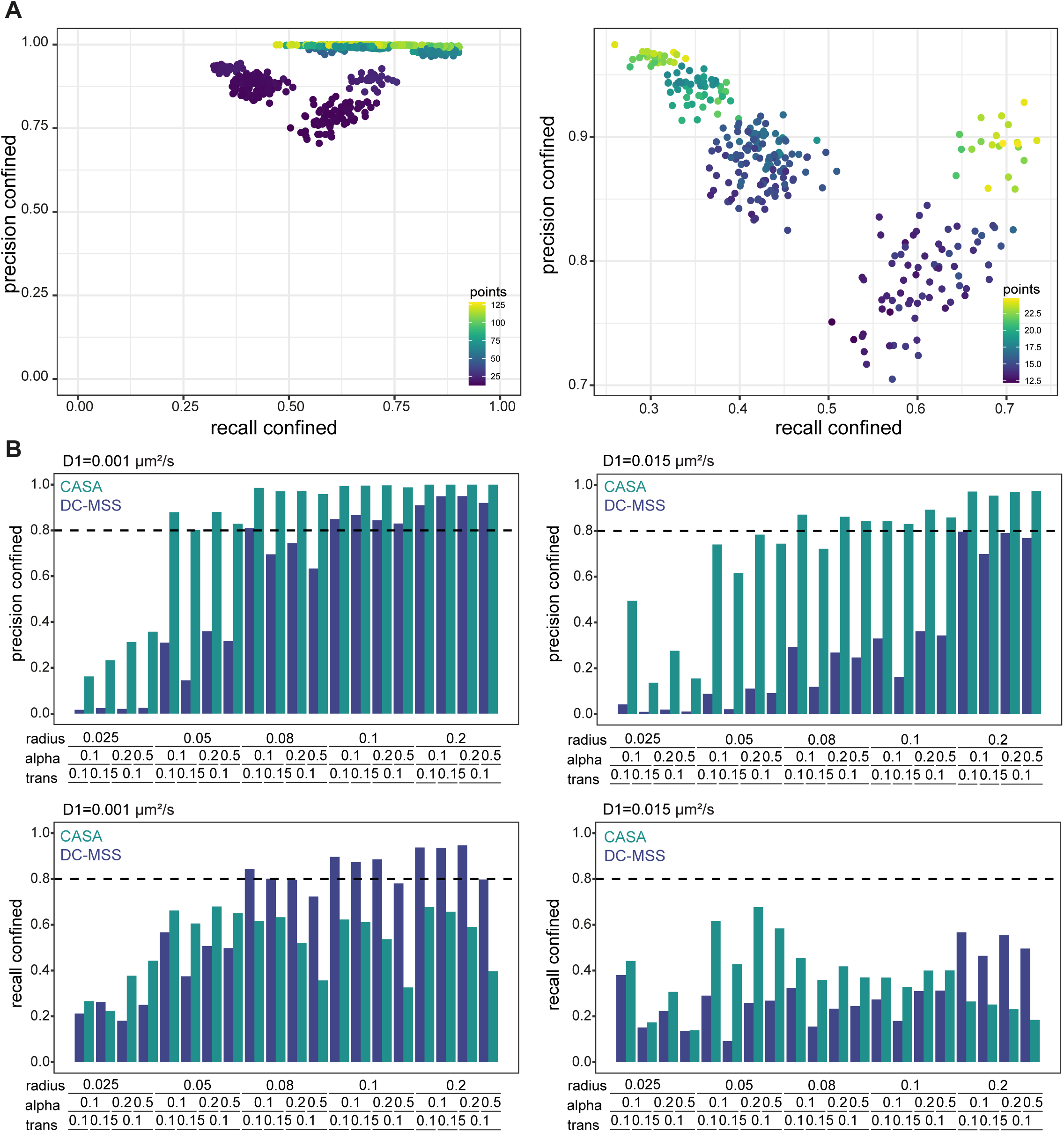
Classifier performance evaluation. **A.** Scatterplots showing the precision and recall performance on the confined class, color-coded by length of the track containing the confined diffusion steps. On the right, zoom-in on trajectories with 20 points or less. **B.** Mean precision and recall performance of CASTA on the confined class compared to DC-MSS across varying ground-truth simulation parameters.

**Supplementary Movie 1 Effect of 488 nm illumination on mEOS3.2-REM1.2 fluorescence behavior.**

Observation of 3-week-old *Nicotiana benthamiana* leaves transiently transformed with pUb10::mEOS3.2-REM1.2 and image in presence (488 on) and absence of continuous 488 nm light exposure (25 µW). Scale bar indicates 2 µm.

**Supplementary Movie 2 Long-term tracking of mEOS3.2-REM1.2 in cotyledon epidermal cells.**

Five-day-old seedling of *rem1.2*/*rem1.*3/*rem1.4*/pREM1.2::mEOS3.2-REM1.2 was imaged in presence of continuous 488 nm light exposure (25 µW). Scale bar indicates 2 µm.

**Supplementary Table 1:**
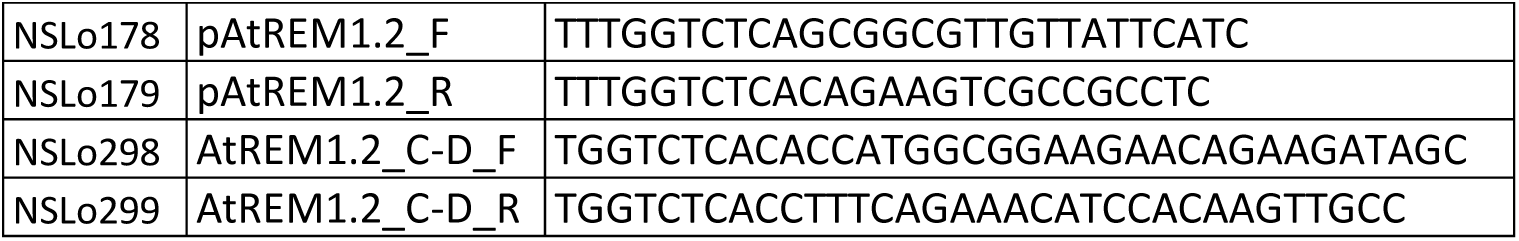
List of primers used in this study.

**Supplementary Table 2:** Feature threshold grid search simulation parameters.

| Parameter | Values | Outside compartment | Inside compartment |
| --- | --- | --- | --- |
| Diffusion coefficient (D) [$\mu\text{m}^2/\text{s}$] | $D_1=0.001$, $D_1=0.005$, $D_1=0.01$, $D_1=0.015$ | $5D_1$, $10D_1$, $25D_1$, $50D_1$ | $1D_1$ |
| Alpha ($\alpha$) | $\alpha=1$ | $1\alpha$ | $0.1\alpha$, $0.15\alpha$ |
| Compartment radius (r) [$\mu\text{m}$] | 0.025, 0.05, 0.1, 0.15 | | |
| Number of compartments (Nc) | 10000 |  |  |
| Transition rate (p) | 0.001, 0.01, 0.1 |  |  |
| Number of particles (Np) | 500 |  |  |

**Supplementary Table 3:** Feature threshold grid search values.

| Feature | Low | Split | High |
| --- | --- | --- | --- |
| Step length | 0.09 | 0.17 | 0.3 |
| Angle | 593 | 740 | 891 |
| KDE | 2 | 31 | 52 |
| Intersections | 5 | 8 | 10 |

**Supplementary Table 4:** Benchmarking simulation parameters.

| Parameter | Values | Outside compartment | Inside Compartment |
| --- | --- | --- | --- |
| Diffusion coefficient (D) [$\mu\text{m}^2/\text{s}$] | $D_1=0.001$, $D_1=0.005$, $D_1=0.01$, $D_1=0.015$, | $5D_1$, $10D_1$, $25D_1$, $50D_1$ | $1D_1$ |
| Alpha ($\alpha$) | $\alpha=1$ | $1\alpha$ | $0.1\alpha$, $0.15\alpha$ |
| Compartment radius (r) [$\mu\text{m}$] | 0.05, 0.08, 0.1, 0.15, 0.2 | | |
| Number of compartments (N) | 8000, 10000 |  |  |
| Transition rate (p) | 0.001, 0.01, 0.1 |  |  |
| Number of particles | 500 |  |  |

## MATERIAL AND METHODS

### Plant materials and culture

All plants used in this work were Arabidopsis thaliana L. (Heyn.), accession Columbia 0. Seeds were stratified at 4°C for 2 days and germinated on half-strength Murashige and Skoog (½ MS) medium supplemented with sucrose (10 g L-1) and agar (8 g L-1) (Murashige and Skoog, 1962) and grown in chambers under controlled conditions with 70% relative humidity and cycles of 16 h of light (∼150 µE m^-2^ sec^-1^) and 8 h of dark at 21 °C. The Arabidopsis transgenic lines Col-0/pUb::mEOS2-AHA2 (Martinière et al., 2019), Col-0/pPIP2;1::PIP2;1-mEOS2 (Hosy et al., 2015), Col-0/pUb::mEOS2-ROP6 (Platre et al., 2019), and Col-0/pUb Lti6a-mEOS2 (Hosy et al., 2015) were previously described. The *rem1.2/rem1.3/rem1.4/*pREM1.2::mEOS3.2-REM1.2 line was generating by transforming the *rem1.2/rem1.3/rem1.4* T-DNA mutant line (Abel et al., 2021; Jolivet et al., 2023) with pREM1.2::mEOS3.2-REM1.2 plasmid using floral dipping (Clough and Bent, 1998). 3-weeks-old Nicotiana benthamiana leaves were transiently transformed using a solution (OD=0.1) of *Agrobacterium tumefaciens* strain GV3101 carrying corresponding constructs as previously described (Marillonnet et al., 2004).

### Molecular cloning

The REM1.2 coding sequence was amplified using Arabidopsis Col-0 complementary DNA (cDNA) and the native promoter of REM1.2 (pREM1.2) was amplified using Arabidopsis Col-0 genomic DNA (gDNA) as templates and the primers listed in the Supplementary Table 1. Both sequences were inserted into a pUC57-based LVL1 golden gate plasmid (Binder et al., 2014) by blunt-end cloning. The codon-optimized coding sequences of mEOS2 and mEOS3.2 were synthetized as gene fragments by TwistBioscience and cloned into a pUC57-based LVL1 golden gate plasmid (Binder et al., 2014), position B-C, using the type II restriction enzyme BpiI. The resulting constructs were used to generate Xpre2-S (pCAMBIA) binary LVL2 plasmids encoding for REM1.2 N-terminally tagged with mEOS variants under the control of the UBQ10 promoter (pUb) or native promoter (pUb::mEOS2-REM1.2, pUb::mEOS3.2-REM1.2 or pREM1.2::mEOS3.2-REM1.2). All constructs were verified by sequencing. The binary vectors were transferred into *Agrobacterium tumefaciens* strain GV3101 for flower dip transformation of Arabidopsis (Clough and Bent, 1998) or transient transformation of *Nicotiana benthamiana* leaves.

### Variable angle-total internal reflection microscopy (VA-TIRFM)

VA-TIRM imaging was operated on a custom-build TIRFM set-up (Rohr et al., 2024), equipped with an 100x objective NA 1.49 (Zeiss, 421190-9800-000), four laser lines (405, 488, 561 and 642 nm), a polychromatic modulator (AOTF, AA OPTO-ELECTRONIC) and a sCMOS camera (Hamamatsu Photonics, ORCA-Flash4.0 V2). To ensure that the target laser power value always corresponds to the actual irradiance in the sample plane, the power of each laser line was calibrated before each experiment using a PM100D detector (Thorlabs). Arabidopsis five-days-old seedlings or leaf discs (4 mm in diameter) of 3-weeks-old N. benthamiana leaves (28 to 34 hours post agrobacterium infiltration) were mounted between two coverslips (Epredia 24×50 mm #1) in liquid ½ MS medium and delicately placed on the specimen stage without additional weight. Images were acquired at 50 Hz frame rate, using 1000 µW 561nm laser power, with or without 1-3µW 405 nm laser power and with or without 1-30 µW 488 nm laser power. The emitted light was filtered using 568 LP Edge Basic Long-pass Filter, 584/40 ET Band-pass filters (AHF analysentechnik AG) and recorded from a 51.2 x 51.2 µm region of interest, 100 nm pixel size.

### Single-particle tracking photoactivated localization microscopy (spt-PALM)

To ensure bona-fide single-molecule tracking we analysed frames with relatively low molecule density (ca. 0.1 – 1 molecule per µm^2^). We used the plugin TrackMate 7 (Ershov et al., 2022) in Fiji (Schindelin et al., 2012) to reconstruct single-molecule trajectories. Single particles were segmented frame-by-frame by applying a Laplacian of Gaussian (LoG) filter and estimated particle size of 0.3 μm. Individual single particles were localized with sub-pixel resolution using a built-in quadratic fitting scheme. Single-particle trajectories were reconstructed using a simple linear assignment problem (Jaqaman et al., 2008) with a maximal linking distance of 0.4 μm and a 4 frames-gap-closing maximum. The coordinates of the single-particle trajectories were then further analysed using a custom-made python script to calculate the mean square displacement (MSD) and apparent diffusion coefficient (D) in batch mode. Only tracks with a minimum of five localizations were kept for analysis. The MSD and diffusion coefficient was calculated based on the first four time points of each trajectory as previously described (Saxton and Jacobson, 2003).

### Core Classifier training

A Hidden Markov Model (HMM) was trained on N = 30 000 simulated ground truth tracjectories. For this, we used the simulation framework developed by AnDi (Muñoz-Gil et al., 2025), which allows to simulate single tracks, that are stochastically entering or leaving confined zones. From the raw track data (sequences of x,y,t coordinate pairs), we calculated the step length, MSD and logD in a sliding window of length ten along each track and independently standardized each feature sequence using the StandardScaler implementation from sklearn.preprocessing. The standardized features then served as input for the HMM (Supplemental Figure 10 A, B). The model was implemented using the hmmlearn package (https://github.com/hmmlearn/hmmlearn), using the Viterbi algorithm option to find the most likely sequence of hidden states given the input features, and was run for 100 iterations from ten random initial states, selecting the best performing model after training.

### Diffusional signature metrics

For each trajectory time point, a suite of four quantitative diffusional features is computed to characterize local motion dynamics in a sliding window of ten consecutive steps. First, since spatially arrested molecules tend to have shorter track motion steps between consecutive time points, we calculate the step length in a sliding window (Schirripa Spagnolo & Luin, 2024). Secondly, we considered that spatially arrested molecules exhibit smaller angles between successive localization steps compared to diffusive molecules (Schirripa Spagnolo & Luin, 2024). Another widely-used metric to assess local density heterogeneities of single-particle localizations is 2-dimensional Kernel density estimation (KDE) (Wallis et al., 2023). Thus, as a third criterion, track-wise KDEs are computed. To obtain a binary classification based on KDE, KDE values are categorized into equal bins and inverted. Fourth, we considered that when a molecule enters a more spatially arrested state, it is more prone to intersect its previously travelled path at these locations. We therefore computed the self-intersections of each track using a sliding window (i.e., for time step n=1, we computed the intersections with n=4, 5, 6 and 7; n=2 with n=5, 6, 7 and 8, etc.). To estimate the parameter value that maximizes the separation between normal diffusing and spatially arrested time steps for each metric, we optimized the classification thresholds for the four computed track features. We generated a set of ground truth particle tracks by simulating single-molecule diffusion across a grid search of simulation parameters (Supplementary Table 2). On this dataset, we tested the performance of the classifier by varying candidate threshold values for each feature: The mean value of the positive and negative ground truth classes, and the value providing the best separation between the two class distributions (Supplementary Table 3, Supplementary Figure 10 C). For each classification run, we computed the False Positive Rate (FPR) and True Positive Rate (TPR) and subsequently calculated the Area Under the Curve (AUC) score for each feature threshold value (Supplementary Figure 10 D).

### Simulations and benchmarking

To simulate ground truth data, we used the simulation framework developed by (Muñoz-Gil et al., 2025). For all simulations we generated 500 particles with a track length of 200 timepoints and a frame rate of 0.1s. The simulated field of view was 128×128px^2^, with a pixel size of 100 nm. Further, to account for the inherent localization imprecision of optical detection systems, we used a localization error of sigma = 12 nm for all our simulations. Based on *in vivo* observations, we used a 5- to 50-fold higher diffusion rate outside of transiently confined areas compared to confined zones (Low-Nam et al., 2011; Suzuki et al., 2007). Similarly, for the anomalous diffusion exponent α, we used a range below 1 for transiently confined segments. To simulate particles stochastically entering and leaving confinement zones, we used a range of different transition probabilities (p) as well as different confinement radii (r) (Supplementary Table 3, 4). To benchmark the performance of CASTA, we generated a set of 480 simulations with unique parameter combinations (Supplementary Table 4). Mean and standard deviation of three independent simulations per combination were used to calculate precision and recall. To compare CASTAs performance to DC-MSS, a smaller set of 40 different simulations, with N = 8 000 compartments and parameter combinations as indicated in (Supplementary Figure 11 C) was generated. In order to compare the 5 states of DC-MSS (0, 1, 2, 3, 4) to the 2 states of CASTA, we re-assigned the following states: 0, 1 = diffusive, 2, 3, 4 = arrested.

### Statistical analysis

The number of independent experiments and the number of individual cells analysed per condition and collected across these experiments are indicated in each figure legends. The statical tests used are reported in the figure legends and have been performed using R or GraphPad Prism.

## REFERENCES

Asghar, S., Ni, R., & Volpe, G. (2025). U-Net 3+ for anomalous diffusion analysis enhanced with mixture estimates (U-AnD-ME) in particle-tracking data. Journal of Physics: Photonics, 7(4), 045005. 10.1088/2515-7647/adf9aa

Bayle, V., Fiche, J.-B., Burny, C., Platre, M. P., Nollmann, M., Martinière, A., & Jaillais, Y. (2021). Single-particle tracking photoactivated localization microscopy of membrane proteins in living plant tissues. Nature Protocols, 16(3), 1600–1628. 10.1038/s41596-020-00471-4

Betzig, E., Patterson, G. H., Sougrat, R., Lindwasser, O. W., Olenych, S., Bonifacino, J. S., Davidson, M. W., Lippincott-Schwartz, J., & Hess, H. F. (2006). Imaging intracellular fluorescent proteins at nanometer resolution. Science, 313(5793), 1642–1645. 10.1126/science.1127344

Bücherl, C. A., Jarsch, I. K., Schudoma, C., Segonzac, C., Mbengue, M., Robatzek, S., MacLean, D., Ott, T., & Zipfel, C. (2017). Plant immune and growth receptors share common signalling components but localise to distinct plasma membrane nanodomains. Elife, 6. 10.7554/eLife.25114

Das, R., Cairo, C. W., & Coombs, D. (2009). A Hidden Markov Model for Single Particle Tracks Quantifies Dynamic Interactions between LFA-1 and the Actin Cytoskeleton. PLOS Computational Biology, 5(11), e1000556. 10.1371/journal.pcbi.1000556

Daumas, F., Destainville, N., Millot, C., Lopez, A., Dean, D., & Salomé, L. (2003). Confined Diffusion Without Fences of a G-Protein-Coupled Receptor as Revealed by Single Particle Tracking. Biophysical Journal, 84(1), 356–366. 10.1016/S0006-3495(03)74856-5

De Zitter, E., Thédié, D., Mönkemöller, V., Hugelier, S., Beaudouin, J., Adam, V., Byrdin, M., Van Meervelt, L., Dedecker, P., & Bourgeois, D. (2019). Mechanistic investigation of mEos4b reveals a strategy to reduce track interruptions in sptPALM. Nat Methods, 16(8), 707–710. 10.1038/s41592-019-0462-3

Demir, F., Horntrich, C., Blachutzik, J. O., Scherzer, S., Reinders, Y., Kierszniowska, S., Schulze, W. X., Harms, G. S., Hedrich, R., Geiger, D., & Kreuzer, I. (2013). Arabidopsis nanodomain-delimited ABA signaling pathway regulates the anion channel SLAH3. Proc Natl Acad Sci U S A, 110(20), 8296–8301. 10.1073/pnas.1211667110

Dosset, P., Rassam, P., Fernandez, L., Espenel, C., Rubinstein, E., Margeat, E., & Milhiet, P.-E. (2016). Automatic detection of diffusion modes within biological membranes using back-propagation neural network. BMC Bioinformatics, 17(1), 197. 10.1186/s12859-016-1064-z

D’Este, E., Lukinavičius, G., Lincoln, R., Opazo, F., & Fornasiero, E. F. (2024). Advancing cell biology with nanoscale fluorescence imaging: essential practical considerations. Trends Cell Biol, 34(8), 671–684. 10.1016/j.tcb.2023.12.001

Granik, N., Weiss, L. E., Nehme, E., Levin, M., Chein, M., Perlson, E., Roichman, Y., & Shechtman, Y. (2019). Single-Particle Diffusion Characterization by Deep Learning. Biophysical Journal, 117(2), 185–192. 10.1016/j.bpj.2019.06.015

Gronnier, J., Crowet, J.-M., Habenstein, B., Nasir, M. N., Bayle, V., Hosy, E., Platre, M. P., Gouguet, P., Raffaele, S., Martinez, D., Grelard, A., Loquet, A., Simon-Plas, F., Gerbeau-Pissot, P., Der, C., Bayer, E. M., Jaillais, Y., Deleu, M., Germain, V.,…Mongrand, S. (2017). Structural basis for plant plasma membrane protein dynamics and organization into functional nanodomains. Elife, 6, e26404. 10.7554/eLife.26404

Gronnier, J., Franck, C. M., Stegmann, M., DeFalco, T. A., Abarca, A., von Arx, M., Dünser, K., Lin, W., Yang, Z., Kleine-Vehn, J., Ringli, C., & Zipfel, C. (2022). Regulation of immune receptor kinase plasma membrane nanoscale organization by a plant peptide hormone and its receptors. Elife, 11, e74162. 10.7554/eLife.74162

Hosy, E., Martinière, A., Choquet, D., Maurel, C., & Luu, D.-T. (2015). Super-Resolved and Dynamic Imaging of Membrane Proteins in Plant Cells Reveal Contrasting Kinetic Profiles and Multiple Confinement Mechanisms. Molecular Plant, 8(2), 339–342. 10.1016/j.molp.2014.10.006

Jaillais, Y., Bayer, E., Bergmann, D. C., Botella, M. A., Boutté, Y., Bozkurt, T. O., Caillaud, M. C., Germain, V., Grossmann, G., Heilmann, I., Hemsley, P. A., Kirchhelle, C., Martinière, A., Miao, Y., Mongrand, S., Müller, S., Noack, L. C., Oda, Y., Ott, T.,…Gronnier, J. (2024). Guidelines for naming and studying plasma membrane domains in plants. Nat Plants, 10(8), 1172–1183. 10.1038/s41477-024-01742-8

Jarsch, I. K., Konrad, S. S., Stratil, T. F., Urbanus, S. L., Szymanski, W., Braun, P., Braun, K. H., & Ott, T. (2014). Plasma Membranes Are Subcompartmentalized into a Plethora of Coexisting and Diverse Microdomains in Arabidopsis and Nicotiana benthamiana. Plant Cell, 26(4), 1698–1711. 10.1105/tpc.114.124446

Jolivet, M. D., Deroubaix, A. F., Boudsocq, M., Abel, N. B., Rocher, M., Robbe, T., Wattelet-Boyer, V., Huard, J., Lefebvre, D., Lu, Y. J., Day, B., Saias, G., Ahmed, J., Cotelle, V., Giovinazzo, N., Gallois, J. L., Yamaji, Y., German-Retana, S., Gronnier, J.,…Germain, V. (2025). Interdependence of plasma membrane nanoscale dynamics of a kinase and its cognate substrate underlies Arabidopsis response to viral infection. Elife, 12. 10.7554/eLife.90309

Kleine-Vehn, J., Wabnik, K., Martinière, A., Łangowski, Ł., Willig, K., Naramoto, S., Leitner, J., Tanaka, H., Jakobs, S., Robert, S., Luschnig, C., Govaerts, W., W Hell, S., Runions, J., & Friml, J. (2011). Recycling, clustering, and endocytosis jointly maintain PIN auxin carrier polarity at the plasma membrane. Molecular Systems Biology, 7(1), MSB201172. 10.1038/msb.2011.72

Konopka, C. A., & Bednarek, S. Y. (2008). Variable-angle epifluorescence microscopy: a new way to look at protein dynamics in the plant cell cortex. The Plant Journal, 53(1), 186–196. 10.1111/j.1365-313X.2007.03306.x

Kowalek, P., Loch-Olszewska, H., & Szwabiński, J. (2019). Classification of diffusion modes in single-particle tracking data: Feature-based versus deep-learning approach. Physical Review E, 100(3), 032410. 10.1103/PhysRevE.100.032410

Kusumi, A., Tsunoyama, T. A., Hirosawa, K. M., Kasai, R. S., & Fujiwara, T. K. (2014). Tracking single molecules at work in living cells. Nature Chemical Biology, 10(7), 524–532. 10.1038/nchembio.1558

Lelek, M., Gyparaki, M. T., Beliu, G., Schueder, F., Griffié, J., Manley, S., Jungmann, R., Sauer, M., Lakadamyali, M., & Zimmer, C. (2021). Single-molecule localization microscopy. Nature Reviews Methods Primers, 1(1), 39. 10.1038/s43586-021-00038-x

Liu, H., Dong, P., Ioannou, M. S., Li, L., Shea, J., Pasolli, H. A., Grimm, J. B., Rivlin, P. K., Lavis, L. D., Koyama, M., & Liu, Z. (2018). Visualizing long-term single-molecule dynamics in vivo by stochastic protein labeling. Proceedings of the National Academy of Sciences, 115(2), 343–348. doi:10.1073/pnas.1713895115

Low-Nam, S. T., Lidke, K. A., Cutler, P. J., Roovers, R. C., van Bergen en Henegouwen, P. M., Wilson, B. S., & Lidke, D. S. (2011). ErbB1 dimerization is promoted by domain co-confinement and stabilized by ligand binding. Nat Struct Mol Biol, 18(11), 1244–1249. 10.1038/nsmb.2135

Ma, Z., Sun, Y., Zhu, X., Yang, L., Chen, X., & Miao, Y. (2022). Membrane nanodomains modulate formin condensation for actin remodeling in Arabidopsis innate immune responses. Plant Cell, 34(1), 374–394. 10.1093/plcell/koab261

Magdziarz, M., Weron, A., Burnecki, K., & Klafter, J. (2009). Fractional Brownian Motion Versus the Continuous-Time Random Walk: A Simple Test for Subdiffusive Dynamics. Physical Review Letters, 103(18), 180602. 10.1103/PhysRevLett.103.180602

Manley, S., Gillette, J. M., Patterson, G. H., Shroff, H., Hess, H. F., Betzig, E., & Lippincott-Schwartz, J. (2008). High-density mapping of single-molecule trajectories with photoactivated localization microscopy. Nature Methods, 5(2), 155–157. 10.1038/nmeth.1176

Matsuda, Y., Hanasaki, I., Iwao, R., Yamaguchi, H., & Niimi, T. (2018). Estimation of diffusive states from single-particle trajectory in heterogeneous medium using machine-learning methods [10.1039/C8CP02566E]. Physical Chemistry Chemical Physics, 20(37), 24099–24108. 10.1039/C8CP02566E

McKinney, S. A., Murphy, C. S., Hazelwood, K. L., Davidson, M. W., & Looger, L. L. (2009). A bright and photostable photoconvertible fluorescent protein. Nature Methods, 6(2), 131–133. 10.1038/nmeth.1296

Michalet, X., Pinaud, F. F., Bentolila, L. A., Tsay, J. M., Doose, S., Li, J. J., Sundaresan, G., Wu, A. M., Gambhir, S. S., & Weiss, S. (2005). Quantum Dots for Live Cells, in Vivo Imaging, and Diagnostics. Science, 307(5709), 538–544. doi:10.1126/science.1104274

Muñoz-Gil, G., Bachimanchi, H., Pineda, J., Midtvedt, B., Fernández-Fernández, G., Requena, B., Ahsini, Y., Asghar, S., Bae, J., Barrantes, F. J., Bender, S. W. B., Cabriel, C., Conejero, J. A., Escoto, M., Feng, X., Haidari, R., Hatzakis, N. S., Huang, Z., Izeddin, I.,…Manzo, C. (2025). Quantitative evaluation of methods to analyze motion changes in single-particle experiments. Nature Communications, 16(1), 6749. 10.1038/s41467-025-61949-x

Muñoz-Gil, G., Garcia-March, M. A., Manzo, C., Martín-Guerrero, J. D., & Lewenstein, M. (2020). Single trajectory characterization via machine learning. New Journal of Physics, 22(1), 013010. 10.1088/1367-2630/ab6065

Muñoz-Gil, G., Volpe, G., Garcia-March, M. A., Aghion, E., Argun, A., Hong, C. B., Bland, T., Bo, S., Conejero, J. A., Firbas, N., Garibo i Orts, Ò., Gentili, A., Huang, Z., Jeon, J.-H., Kabbech, H., Kim, Y., Kowalek, P., Krapf, D., Loch-Olszewska, H.,… Manzo, C. (2021). Objective comparison of methods to decode anomalous diffusion. Nature Communications, 12(1), 6253. 10.1038/s41467-021-26320-w

Nguyen, T. D., Chen, Y.-I., Chen, L. H., & Yeh, H.-C. (2023). Recent Advances in Single-Molecule Tracking and Imaging Techniques. Annual Review of Analytical Chemistry, 16(Volume 16, 2023), 253-284. 10.1146/annurev-anchem-091922-073057

Ott, M., Shai, Y., & Haran, G. (2013). Single-Particle Tracking Reveals Switching of the HIV Fusion Peptide between Two Diffusive Modes in Membranes. The Journal of Physical Chemistry B, 117(42), 13308–13321. 10.1021/jp4039418

Perraki, A., Gronnier, J., Gouguet, P., Boudsocq, M., Deroubaix, A.-F., Simon, V., German-Retana, S., Legrand, A., Habenstein, B., Zipfel, C., Bayer, E., Mongrand, S., & Germain, V. (2018). REM1.3’s phospho-status defines its plasma membrane nanodomain organization and activity in restricting PVX cell-to-cell movement. PLOS Pathogens, 14(11), e1007378. 10.1371/journal.ppat.1007378

Persson, F., Lindén, M., Unoson, C., & Elf, J. (2013). Extracting intracellular diffusive states and transition rates from single-molecule tracking data. Nature Methods, 10(3), 265–269. 10.1038/nmeth.2367

Platre, M. P., Bayle, V., Armengot, L., Bareille, J., Marquès-Bueno, M. D. M., Creff, A., Maneta-Peyret, L., Fiche, J. B., Nollmann, M., Miège, C., Moreau, P., Martinière, A., & Jaillais, Y. (2019). Developmental control of plant Rho GTPase nano-organization by the lipid phosphatidylserine. Science, 364(6435), 57–62. 10.1126/science.aav9959

Rabiner, L. R. (2002). A tutorial on hidden Markov models and selected applications in speech recognition. Proceedings of the IEEE, 77(2), 257–286.

Raffaele, S., Bayer, E., Lafarge, D., Cluzet, S., German Retana, S., Boubekeur, T., Leborgne-Castel, N., Carde, J. P., Lherminier, J., Noirot, E., Satiat-Jeunemaître, B., Laroche-Traineau, J., Moreau, P., Ott, T., Maule, A. J., Reymond, P., Simon-Plas, F., Farmer, E. E., Bessoule, J. J., & Mongrand, S. (2009). Remorin, a solanaceae protein resident in membrane rafts and plasmodesmata, impairs potato virus X movement. Plant Cell, 21(5), 1541–1555. 10.1105/tpc.108.064279

Rodriguez, E. A., Campbell, R. E., Lin, J. Y., Lin, M. Z., Miyawaki, A., Palmer, A. E., Shu, X., Zhang, J., & Tsien, R. Y. (2017). The Growing and Glowing Toolbox of Fluorescent and Photoactive Proteins. Trends in Biochemical Sciences, 42(2), 111–129. 10.1016/j.tibs.2016.09.010

Rohr, L., Ehinger, A., Rausch, L., Glöckner Burmeister, N., Meixner, A. J., Gronnier, J., Harter, K., Kemmerling, B., & zur Oven-Krockhaus, S. (2024). OneFlowTraX: a user-friendly software for super-resolution analysis of single-molecule dynamics and nanoscale organization [Technology and Code]. Frontiers in Plant Science, Volume 15 - 2024. 10.3389/fpls.2024.1358935

Rohr, L., Rausch, L., Harter, K., & zur Oven-Krockhaus, S. (2024). Contrasting Effects of Cytoskeleton Disruption on Plasma Membrane Receptor Dynamics: Insights from Single-Molecule Analyses. bioRxiv, 2024.2009.2009.612020. 10.1101/2024.09.09.612020

Schirripa Spagnolo, C., & Luin, S. (2024). Trajectory Analysis in Single-Particle Tracking: From Mean Squared Displacement to Machine Learning Approaches. Int J Mol Sci, 25(16). 10.3390/ijms25168660

Shroff, H., Galbraith, C. G., Galbraith, J. A., White, H., Gillette, J., Olenych, S., Davidson, M. W., & Betzig, E. (2007). Dual-color superresolution imaging of genetically expressed probes within individual adhesion complexes. Proceedings of the National Academy of Sciences, 104(51), 20308–20313. doi:10.1073/pnas.0710517105

Smokvarska, M., Bayle, V., Maneta-Peyret, L., Fouillen, L., Poitout, A., Dongois, A., Fiche, J.-B., Gronnier, J., Garcia, J., Höfte, H., Nolmann, M., Zipfel, C., Maurel, C., Moreau, P., Jaillais, Y., & Martiniere, A. (2023). The receptor kinase FERONIA regulates phosphatidylserine localization at the cell surface to modulate ROP signaling. Science Advances, 9(14), eadd4791. doi:10.1126/sciadv.add4791

Smokvarska, M., Francis, C., Platre, M. P., Fiche, J. B., Alcon, C., Dumont, X., Nacry, P., Bayle, V., Nollmann, M., Maurel, C., Jaillais, Y., & Martiniere, A. (2020). A Plasma Membrane Nanodomain Ensures Signal Specificity during Osmotic Signaling in Plants. Curr Biol, 30(23), 4654–4664.e4654. 10.1016/j.cub.2020.09.013

Suzuki, K. G., Fujiwara, T. K., Sanematsu, F., Iino, R., Edidin, M., & Kusumi, A. (2007). GPI-anchored receptor clusters transiently recruit Lyn and G alpha for temporary cluster immobilization and Lyn activation: single-molecule tracking study 1. J Cell Biol, 177(4), 717–730. 10.1083/jcb.200609174

Tabei, S. M. A., Burov, S., Kim, H. Y., Kuznetsov, A., Huynh, T., Jureller, J., Philipson, L. H., Dinner, A. R., & Scherer, N. F. (2013). Intracellular transport of insulin granules is a subordinated random walk. Proceedings of the National Academy of Sciences, 110(13), 4911–4916. doi:10.1073/pnas.1221962110

Thapa, S., Lomholt, M. A., Krog, J., Cherstvy, A. G., & Metzler, R. (2018). Bayesian analysis of single-particle tracking data using the nested-sampling algorithm: maximum-likelihood model selection applied to stochastic-diffusivity data [10.1039/C8CP04043E]. Physical Chemistry Chemical Physics, 20(46), 29018–29037. 10.1039/C8CP04043E

Tokunaga, M., Imamoto, N., & Sakata-Sogawa, K. (2008). Highly inclined thin illumination enables clear single-molecule imaging in cells. Nat Methods, 5(2), 159–161. 10.1038/nmeth1171

Vega, A. R., Freeman, S. A., Grinstein, S., & Jaqaman, K. (2018). Multistep Track Segmentation and Motion Classification for Transient Mobility Analysis. Biophys J, 114(5), 1018–1025. 10.1016/j.bpj.2018.01.012

von Arx, M., Jolivet, M.-D., Biermann, D., Gabani, V., Andrews, S. S., Zipfel, C., & Gronnier, J. (2026). Plasma membrane nanoscale dynamics of Arabidopsis leucine-rich repeat receptor kinase complexes. bioRxiv. 10.64898/2026.03.05.709869

Wallis, T. P., Jiang, A., Young, K., Hou, H., Kudo, K., McCann, A. J., Durisic, N., Joensuu, M., Oelz, D., Nguyen, H., Gormal, R. S., & Meunier, F. A. (2023). Super-resolved trajectory-derived nanoclustering analysis using spatiotemporal indexing. Nature Communications, 14(1), 3353. 10.1038/s41467-023-38866-y

Wang, S., Moffitt, J. R., Dempsey, G. T., Xie, X. S., & Zhuang, X. (2014). Characterization and development of photoactivatable fluorescent proteins for single-molecule–based superresolution imaging. Proceedings of the National Academy of Sciences, 111(23), 8452–8457. doi:10.1073/pnas.1406593111

Weigel, A. V., Simon, B., Tamkun, M. M., & Krapf, D. (2011). Ergodic and nonergodic processes coexist in the plasma membrane as observed by single-molecule tracking. Proc Natl Acad Sci U S A, 108(16), 6438–6443. 10.1073/pnas.1016325108

Zhang, M., Chang, H., Zhang, Y., Yu, J., Wu, L., Ji, W., Chen, J., Liu, B., Lu, J., Liu, Y., Zhang, J., Xu, P., & Xu, T. (2012). Rational design of true monomeric and bright photoactivatable fluorescent proteins. Nature Methods, 9(7), 727–729. 10.1038/nmeth.2021

